# Neural System and Receptor Diversity in the ctenophore *Beroe abyssicola*

**DOI:** 10.1101/419218

**Authors:** Tigran P. Norekian, Leonid L. Moroz

## Abstract

Although, neuro-sensory systems might evolve independently in ctenophores, very little is known about their neural organization. Most of the ctenophores are pelagic and deep-water species and cannot be bred in the laboratory. Thus, it is not surprising that neuroanatomical data are available for only one genus within the group - *Pleurobrachia.* Here, using immunohistochemistry and scanning electron microscopy, we describe the organization of two distinct neural subsystems (subepithelial and mesogleal) and the structure of different receptor types in the comb jelly *Beroe abyssicola -* the voracious predator from North Pacific. A complex subepithelial neural network of *Beroe,* with five receptor types, covers the entire body surface and expands deeply into the pharynx. Three types of mesogleal neurons are comparable to the cydippid *Pleurobrachia*. The predatory lifestyle of *Beroe* is supported by the extensive development of ciliated and muscular structures including the presence of giant muscles and feeding macrocilia. The obtained neuroanatomy atlas provides unique examples of lineage-specific innovations within these enigmatic marine animals, and remarkable complexity of sensory and effector systems in this clade of basal Metazoa.

**Graphical Abstract:** Although, neuro-sensory systems might evolve independently in ctenophores, very little is known about their neuroanatomy. Here, using immunohistochemistry and scanning electron microscopy, we describe the organization of two neural systems and five different receptor types in the comb jelly *Beroe abyssicola -* the voracious predator from North Pacific. The predatory lifestyle of *Beroe* is supported by the extensive development of ciliated, muscular, and nervous systems including the presence of giant muscles and exceptional feeding macrocilia. The obtained neuroanatomy atlas provides unique examples of lineage-specific innovations within this enigmatic group of marine animals.

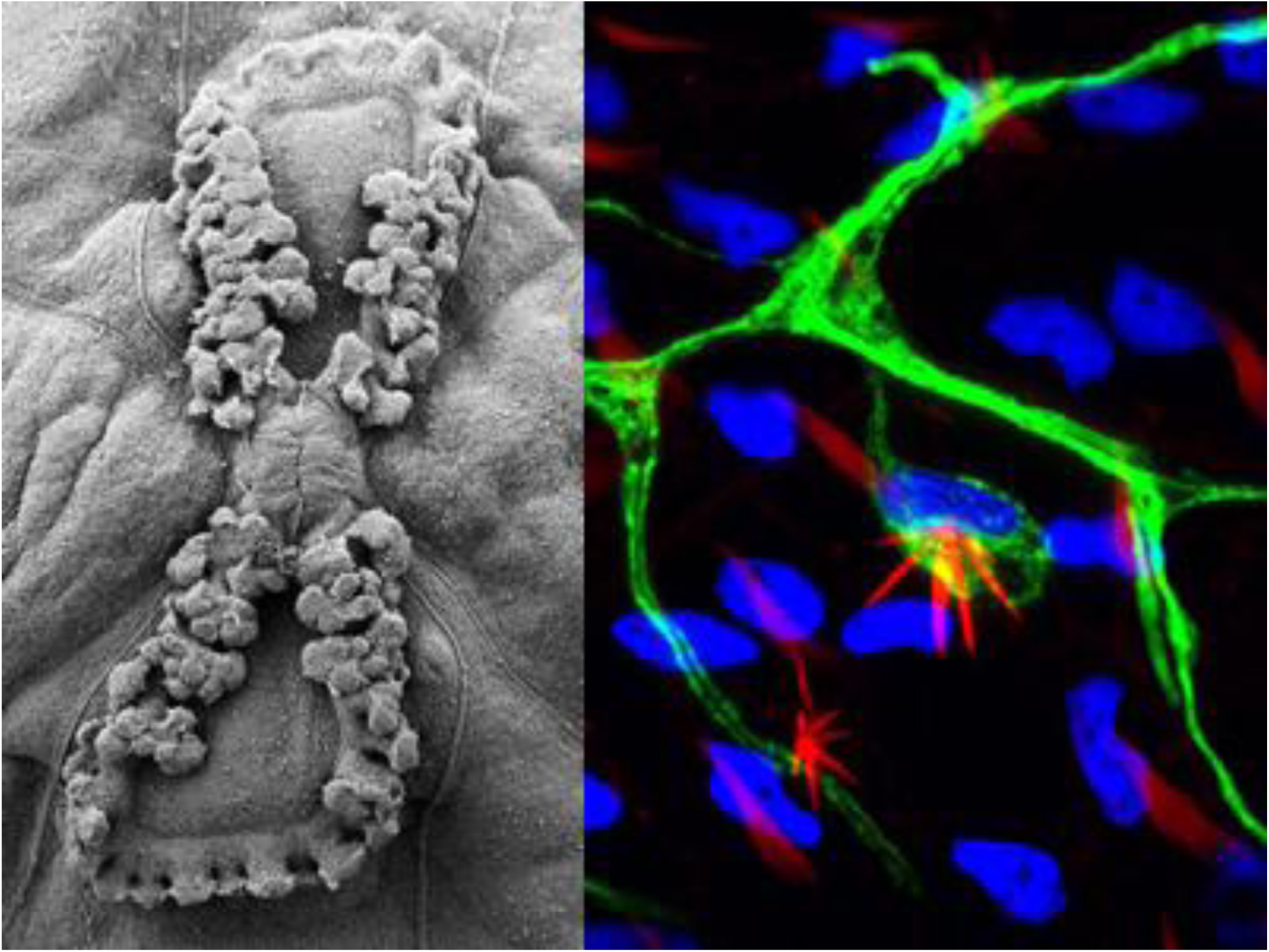

## 1 INTRODUCTION

Ctenophores are one of the basal metazoan lineages comprising of about 200 marine carnivorous species (Tamm, 1982; Kozloff, 1990; Hernandez-Nicaise, 1991). Most of them are pelagic and deep-water organisms, and cannot be bred in the laboratory. Thus, it is not surprising that neuroanatomical data are available for the only one genus within the group *-Pleurobrachia* (Jager et al., 2011; Jager et al., 2013; Norekian and Moroz, 2016; 2018), which genome was also recently sequenced (Moroz et al., 2014). This work, together with genomescale analyses on cnidarians, sponges, and placozoans, provides molecular and phylogenetic evidence that neurons and muscles, as well as mesoderm, might evolve more than once (Moroz, 2009; 2012; Steinmetz et al., 2012; Jorgensen, 2014; Moroz, 2014; 2015; Moroz and Kohn, 2015; 2016). But the field requires broader comparative data including the neuroanatomical mapping of ecologically and morphologically diverse ctenophore species such as *Beroe* (Fig. 1).

**Figure 1.**
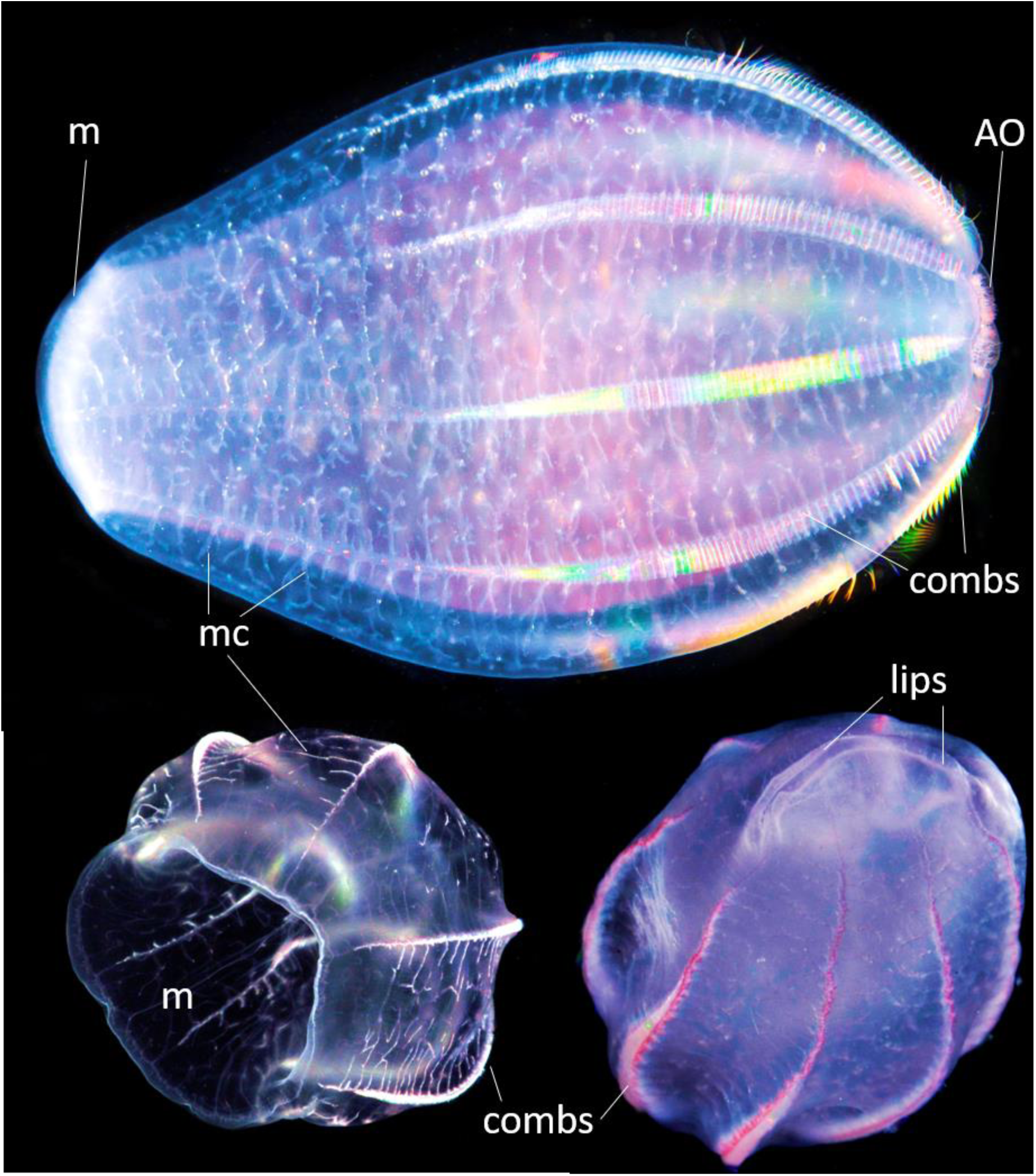
The nontentaculate free swimming ctenophore *Beroe abyssicola;* Mortensen, 1927. *Beroe* has a flattened and highly muscular body with numerous diverticula of meridional canals (me) - a part of the gastrovascular system. *Beroe* constantly swims with the help of eight ciliated comb plates, which show beautiful iridescence under illumination. Combs have one of the largest cilia in the animal kingdom. *Beroe* can actively search and catch its prey *Bolinopsis infundibulum* by the lips, swallowing them with a very wide mouth (m). Swimming and other behaviors can be controlled by the aboral organ (AO) with a gravity sensor. *Beroe* can rapidly change shapes of the body; sizes of animals on the photo are 4-5 cm. See text for further details.

The placement of ctenophores within the animal tree of life is still highly debated (Dunn et al., 2015; Telford et al., 2016; Dunn, 2017; King and Rokas, 2017; Simion et al., 2017), with recent reconstructions supporting the hypothesis that Ctenophores is the sister clade to the rest of Metazoa (Borowiec et al., 2015; Whelan et al., 2015; Halanych et al., 2016; Telford et al., 2016; Arcila et al., 2017; Shen et al., 2017; Whelan et al., 2017). Such scenario means that the extant ctenophores are descendants of the earliest animal lineage, which branched before the subsequent divergence of nerveless sponges and placozoans as well as Eumetazoa (a clade composed of cnidarians and bilaterians). The proposed relationships among basal metazoans provide additional support for the convergent evolution of neurons and synapses (Moroz, 2015; Moroz and Kohn, 2016).

Molecular clock analyses suggest that the modern ctenophore diversity originated approximately 350 million years ago, followed their rapid radiation (Whelan et al., 2017). Consequently, the numerous innovations within the clade are expected and frequently observed (Moroz et al., 2014; Tamm, 2014; Dunn et al., 2015; Kohn et al., 2015; Whelan et al., 2017). For example, the phylogenomic data recognize at least seven major ctenophore groups with Beroidae as the only lineage, which secondarily lost tentacles, possibly, because of switching to different food sources. In fact, both *Beroe* and *Neis* feed on large ctenophores (vs. small planktonic organisms as other comb jellies - see (Harbison et al., 1978)).

The active predatory lifestyle *(Beroe* always swims!) led to a substantial reorganization of the body plan in Beroidae. In addition to the loss of tentacles, there are the extensive development of the gut and very complex hunting behaviors. Upon the contact with the prey, *Beroe* quickly re-orients mouth and engulf whole prey in its gaping lips. The specialized saber-shaped giant cilia (known as macrocilia) help to bite-off pieces of prey (Horridge, 1965a; Swanberg, 1974; Tamm, 1983).

Adaptations to the predation also include specialized reversible epithelial adhesive mechanisms of how to keep the mouth closed without muscular or neuronal activity (Tamm and Tamm, 1991b). Synapses and neurites were identified by electron microscopy at the end of these specialized adhesive cells (Tamm and Tamm, 1991b). But even between two closely related *Beroe* species, there are the substantial differences in the anatomy when the reversible epithelial adhesive ultrastructure was compared (Tamm and Tamm, 1991b). Additional adaptations are giant muscles (Bilbaut et al., 1988), large mitochondria and bioenergetics, critical for active predation, as well as antimicrobial molecules, photoproteins (Markova et al., 2012; Stepanyuk et al., 2013; Burakova et al., 2016); plusdistinct developmental mechanisms (Carre and Sardet, 1984; Carre et al., 1991; Houliston et al., 1993; Rouviere et al., 1994; 1998).

Despite great interest in *Beroe,* previous neuroanatomical studies were focused on distantly related *Pleurobrachia* as the representative of cydippid ctenophores; which has a passive feeding mechanism by capturing small zooplankton with the net of tentacles and tentillae. It is estimated that *Pleurobrachia* and *Beroe* lineages diverged about 250 MYA, around the time of great Permian extinction event (Whelan et al., 2017). Here, we ask whether neural and receptive systems of *Beroe* are comparable to *Pleurobrachia*.

Previous transmission electron microscopy and vital staining with reduced methylene blue indicate the presence of two nerve nets and characteristic synapses in *Beroe* (Hernandez-Nicaise, 1973b; a; c; Hernandez-Nicaise, 1991). Using immunohistochemistry and scanning electron microscopy, we expand our earlier studies on *Pleurobrachia bachei* (Norekian and Moroz, 2016; 2018), and characterize the neural and sensory organization in the *Beroe abyssicola* from North Pacific. The provided neuroanatomical atlas of *Beroe* cell types can be an imperative resource for future molecular characterization of enigmatic neural systems in ctenophores, which is important for the entire field of comparative and evolutionary neuroscience.

## 2 MATERIALS AND METHODS

### 2.1 Animals

Adult specimens of *Beroe abyssicola* (Mortensen, 1927) were collected from the breakwater and held in 1-gallon glass jars in the large tanks with constantly circulating seawater at 10° C. Experiments were carried out at Friday Harbor Laboratories, the University of Washington in the spring-summer seasons of 2012-2018. The details of the immunohistochemical and electron microscopy protocols have been described elsewhere (Norekian and Moroz, 2016; 2018) and we will provide only brief summaries.

### 2.2 Scanning Electron Microscopy (SEM)

Adult, 0.5-5 cm, *Beroe* were fixed in 2.5% glutaraldehyde in 0.2 M phosphate-buffered saline (pH=7.6) for 4 hours at room temperature and washed for 2 hours in 2.5% sodium bicarbonate. Animals then were then dissected into a few smaller pieces. For secondary fixation, we used 2% osmium tetroxide in 1.25% Sodium Bicarbonate for 3 hours at room temperature. The tissue was rinsed several times with distilled water, dehydrated in ethanol and placed in Samdri-790 (Tousimis Research Corporation) for Critical point drying. After the drying process, the tissues were placed on the holding platforms and processed for metal coating on Sputter Coater (SPI Sputter). SEM observations and recordings were done on NeoScope JCM-5000 microscope (JEOL Ltd., Tokyo, Japan).

### 2.3 Immunocytochemistry and phalloidin staining

Adult *Beroe* were fixed overnight in 4% paraformaldehyde in 0.1 M phosphate-buffered saline (PBS) at +4-5° C and washed for 2 hours in PBS. The fixed animals were dissected to get better access to specific organs and preincubated overnight in a blocking solution of 6% goat serum in PBS. The tissues were incubated for 48 hours at +5° C in the primary antibodies diluted in 6% of the goat serum at a final dilution 1:40. We used the rat monoclonal antibody (AbD Serotec Cat# MCA77G, RRID: AB_325003), which recognizes the alpha subunit of tubulin, and specifically binds tyrosylated tubulin (Wehland and Willingham, 1983; Wehland et al., 1983). Following a series of PBS washes for total 6 hours, the tissues were incubated for 12 hours in secondary antibodies - goat anti-rat IgG antibodies (Alexa Fluor 488 conjugated; Molecular Probes, Invitrogen, Cat# A11006, RRID: AB_141373) at a final dilution 1:20.

To label the muscle fibers, we used well-known marker phalloidin (Alexa Fluor 488 and Alexa Fluor 594 phalloidins from Molecular Probes), which binds to F-actin (Wulf et al., 1979). After washing in PBS following the secondary antibody treatment, the tissue was incubated in phalloidin solution in PBS for 4 to 8 hours at a final dilution 1:80 and then washed in several PBS rinses for 6 hours.

To stain nuclei, the tissues were mounted in VECTASHIELD Hard-Set Mounting Medium with DAPI (Cat# H-1500). The preparations were mounted in Fluorescent Mounting Media (KPL) on glass microscope slides to be viewed and photographed using a Nikon Research Microscope Eclipse E800 with Epifluorescence using standard TRITC and FITC filters, BioRad (Radiance 2000 system) Laser Scanning confocal microscope or Nikon C1 Laser Scanning confocal microscope. To test for the specificity of immunostaining either the primary or the secondary antibody were omitted from the procedure. In both cases, no labeling was detected (see other details elsewhere (Norekian and Moroz, 2016; 2018).

### 2.4 Antibody specificity

Rat monoclonal anti-tyrosinated alpha-tubulin antibody is raised against yeast tubulin, clone YL1/2, isotype IgG2a (Serotec Cat # MCA77G; RRID: AB_325003). The antibody is routinely tested in ELISA on tubulin, and the close related ctenophore *Pleurobrachia* is listed in species reactivity (manufacturer’s technical information). The epitope recognized by this antibody appears to be a linear sequence requiring an aromatic residue at the C terminus, with the two adjacent amino acids being negatively charged, represented by Glu-Glu-Tyr in Tyr-Tubulin (manufacturer’s information). As reported by Wehland et al. (Wehland and Willingham, 1983; Wehland et al., 1983)- this rat monoclonal antibody “reacts specifically with the tyrosylated form of brain alpha-tubulin from different species.” They showed that “YL 1/2 reacts with the synthetic peptide Gly-(Glu)3-Gly- (Glu)2-Tyr, corresponding to the carboxyterminal amino acid sequence of tyrosylated alpha-tubulin, but does not react with Gly-(Glu)3-Gly-(Glu)2, the constituent peptide of detyrosinated alpha-tubulin”. The epitope recognized by this antibody has been extensively studied including details about antibody specificity and relevant assays (Wehland and Willingham, 1983; Wehland et al., 1983). Equally important, this specific monoclonal antibody has been used before in two different species of both adult and larval *Pleurobrachia*: the previously obtained staining patterns are very similar to our experiments (Jager et al., 2011; Moroz et al., 2014; Norekian and Moroz, 2016; 2018) as well as successfully tested for 6 other ctenophore species (Norekian and Moroz, unpublished observations).

## 3 RESULTS

### 3.1 Introduction to the *Beroe* anatomy

The pelagic *Beroe abyssicola* and other *Beroe* species, together with the Australian giant *Neis cordigera,* are the only two genera of nontentaculate ctenophores (Kozloff, 1990; Hernandez-Nicaise, 1991), also known as Nuda (Chun, 1879). All Nuda have a significantly different body plan, and are distinguished from other comb jellies, like cydippids (e.g., *Pleurobrachia*) or lobates (e.g., *Mnemiopsis* and *Bolinopsis)* by a complete absence of tentacles, in both juvenile and adult stages.

The North West Pacific, *Beroe abyssicola* (Fig. 1) has a flattened and highly muscular body; it actively swims, searches and catches its prey *Bolinopsis infundibulum* with its very wide mouth. The food is swallowed with the help of strong pharyngeal muscles and macrocilia (see Fig. 16). The huge pharynx extends through most of the body and food is digested within several hours. The pharyngeal and meridional gastrovascular canals form hundreds of branches and run through the entire mesoglea (collagenous, transparent, jelly-like endoskeleton) supplying nutrients and also supporting excretory and reproductive functions.

*Beroe* swims mouth forward using its **eight comb rows** with some of the largest cilia in the animal kingdom (Fig. 2). The scanning electron microscopy (SEM) imaging provided a high-quality assessment of the locomotory system of *Beroe.* Each of eight comb rows consisted of numerous long swim cilia fused together to form a powerful locomotory structure (Fig. 2C). Comb rows covered about 2/3 of the body length stopping short of reaching the aboral organ and extending close to the mouth opening. The individual comb plates were larger in the middle of the row and smoothly narrowed to the end. At the base of each comb, there was a basal cushion standing on the top of mesoglea (Fig. 2C). This cushion consisted of several tall pyramidal cells bearing swim cilia. The **ciliated furrows** coupled each comb row with the aboral organ (Fig. 2D, E, F).

**Figure 2.**
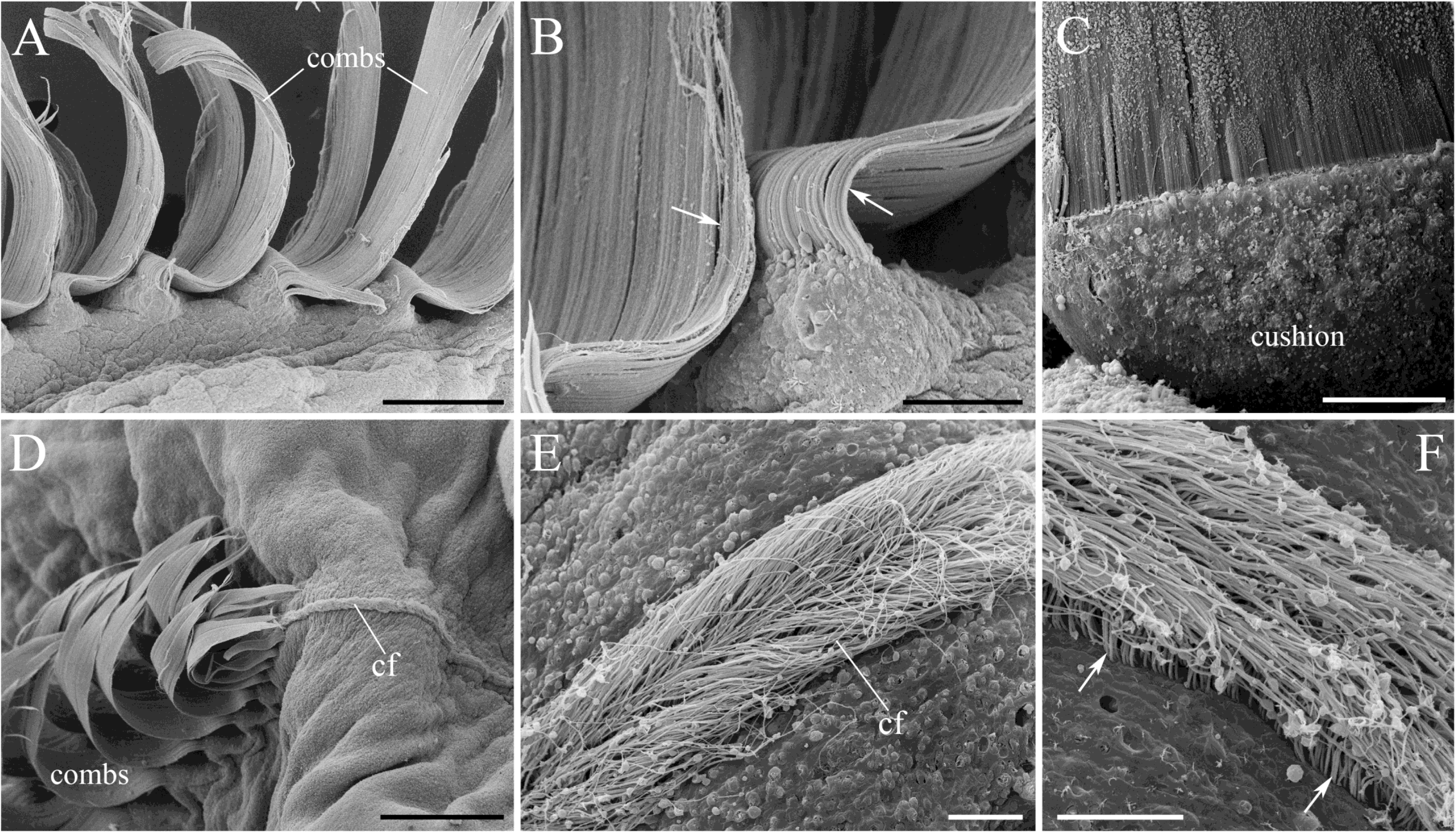
SEM of the combs and ciliated furrows. **A, B**: Comb plates represent a group of long swim cilia fused together (arrows). **C:** A basal cushion, consisting of polster cells that carry swim cilia (arrow), is at the base of each comb. **D:** Each comb row is connected to a ciliated furrow. **E, F:** Ciliated furrow contains a narrow band of densely packed thin cilia (arrows). Scale bars: A - 200 µm; B, C - 40 µm; D - 200 µm; E, F - 10 µm.

The coordination of swimming, gravity sensing and, perhaps, other functions are mediated by **the aboral organ** (Fig. 3A), which is located on the side opposite to the mouth or the oral pole. Two symmetrical **polar fields** are connected to the aboral organ. The term of the aboral organ is preferable to distinguish this analog of “the elementary brain” in ctenophores from the non-homologous apical organ in bilaterian larvae (Nielsen, 2012).

**Figure 3.**
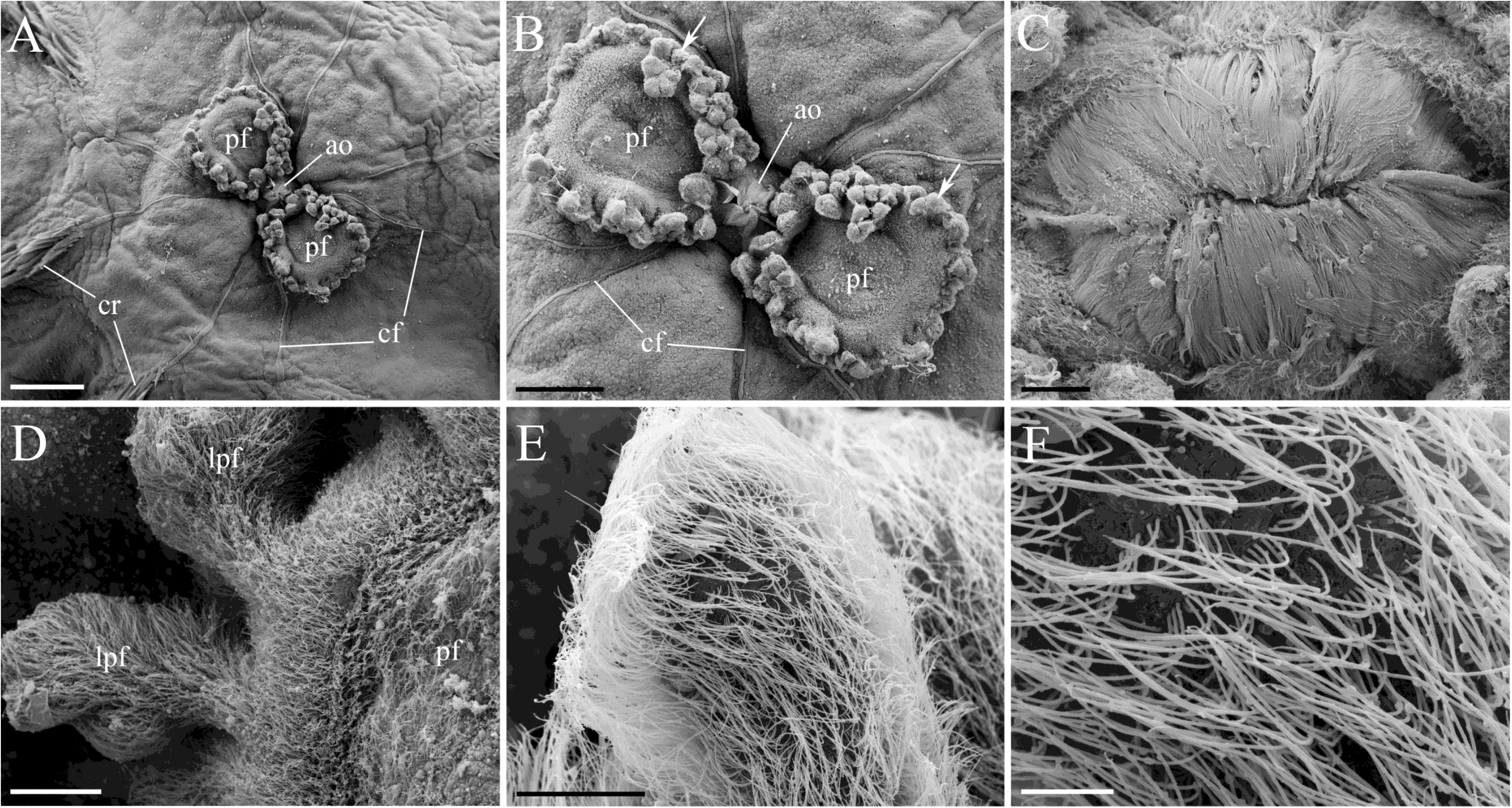
SEM images of the aboral organ and polar fields. **A**: The aboral organ with two polar fields is connected by ciliated furrows to comb rows. **B:** Polar fields in *Beroe* are characterized by a crown of elevated lobes or papillae (arrows). **C:** The aboral organ is covered by a protective dome, which consists of numerous long cilia. **D, E, F:** The lobes of polar fields are covered by long cilia. Abbreviations: *ao* - aboral organ; *pf* - polar field; *Ipf* - lobes of polar field; *cf* - ciliated furrow; *cr* - comb row. Scale bars: A - 500 µm; B - 300 µm; C, D - 50 µm; E - 20 µm; F - 5 µm.

### 3.2 The Aboral Sensory Complex and Polar Fields

SEM imaging in comb jellies is very challenging since these animals are composed >95% of water. However, in highly muscular *Beroe,* the tissues do not go through as much deformation during drying process as in other ctenophores, like *Pleurobrachia* (Norekian and Moroz, 2018).

Eight ciliated furrows (Fig. 3A) represent specialized conductive paths between the aboral organ and the comb rows (Tamm, 1982; Tamm, 2014). The aboral organ itself is covered by a protective dome, which consists of numerous cilia tightly closing the space above the statolith (Fig. 3B, C).

Two polar fields on both sides of the aboral organ are very distinct in *Beroe,* compared to other ctenophores. Each polar field has about 30 large lobes or papillae protruding 100-200 µm up from the epithelial layer and even producing secondary branches (Fig. 3B, C). It creates a 3-dimensional structure, which is more complicated than, for example, very flat polar fields in *Pleurobrachia* or extended curved fields in *Bolinopsis* and *Mnemiopsis*. Long thin cilia densely cover the lobes (Fig. 3D, E, F).

In *Beroe,* as in other ctenophores (Jager et al., 2011; Moroz et al., 2014; Norekian and Moroz, 2016; 2018), the tubulin antibody predominantly labeled neurons and subpopulations of cilia including those covering the entire perimeter of the polar fields and ciliated furrows (Figs. 4A, B; 5A). A diffuse subepithelial neural network was also visualized within the lobes of the polar fields (Fig. 4A).

**Figure 4.**
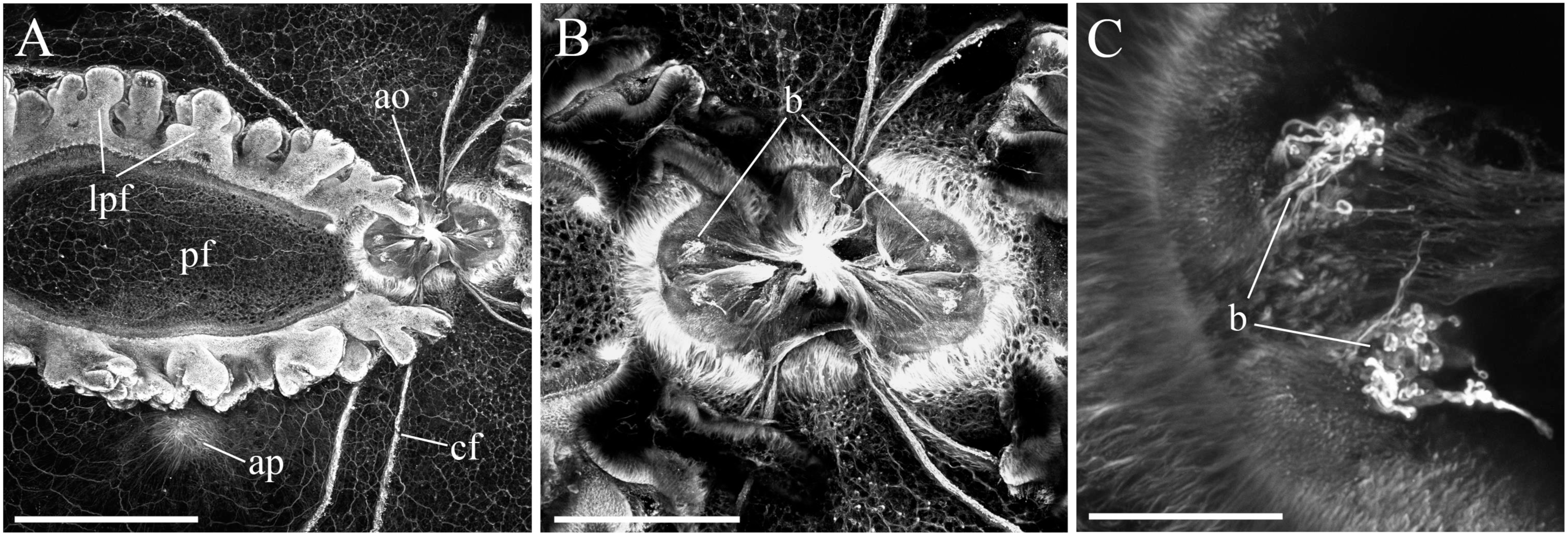
Tubulin IR in the aboral complex. **A**: The aboral organ with one of the polar fields showing the crown of large lobes or papillae and ciliated furrows. Note the diffused network inside and outside of the polar field. **B:** The aboral organ with balancers. **C:** Balancers consist of tight groups of ciliated cells. Abbreviations: *ao* - aboral organ; *pf* - polar field; *Ipf* - lobes of polar field; *cf* - ciliated furrow; *ap* - anal pore; *b* - balancers. Scale bars: A - 500 µm; B - 200 µm; C - 40 µm.

Inside the aboral organ, the tubulin antibody stained four balancers - the structures, which supported the statolith (Figs. 3A, B; 4C) using cilia. Hundreds of compactly packed tubulin-IR cells (2 to 10 µm in diameter) covered the entire floor of the aboral organ (Fig. 5B); some of them revealed elongated processes suggesting their neuron-like nature (Table 1). Phalloidin-stained numerous muscle fibers were attached to the aboral organ from the mesogleal side (Fig. 5A). Contraction of these muscles was responsible for the active defensive withdrawal response of the entire aboral organ and polar fields inside the body of *Beroe.*

**Table 1.** Neuronal cell types identified in *Beroe abyssicola.*

|  | *Neuronal Cell Type* | *Size* | *Location* | *Figure* |
| --- | --- | --- | --- | --- |
| 1. | Neuron-like cells in the aboral organ | 1-7 $\mu\text{m}$ | Aboral organ | Fig. 5 B |
| 2. | Bipolar neurons in the strands of the subepithelial net | 4-7 $\mu\text{m}$ | Subepithelial network | Fig. 7B, C |
| 3. | Tripolar neurons in the nods of the subepithelial net | 4-7 $\mu\text{m}$ | Subepithelial network | Fig. 7B |
| 4. | Neurons located between strands of the subepithelial net | 5-8 $\mu\text{m}$ | Subepithelial network | Figs. 6B, C, D; 7B, C |
| 5. | Neurons located between strands of the subepithelial net with cilia | 5-8 $\mu\text{m}$ | Subepithelial network | Figs. 6C; 11D, E |
| 6. | Bipolar mesogleal neurons | 4-8 $\mu\text{m}$ | Mesoglea | Figs. 8B, C; 9A, B |
| 7. | Multipolar mesogleal neurons | 4-8 $\mu\text{m}$ | Mesoglea | Figs. 8D; 9C, D |
| 8. | Mesogleal neurons projecting a single long neurite | 4-8 $\mu\text{m}$ | Mesoglea | Fig. 8A |

**Figure 5.**
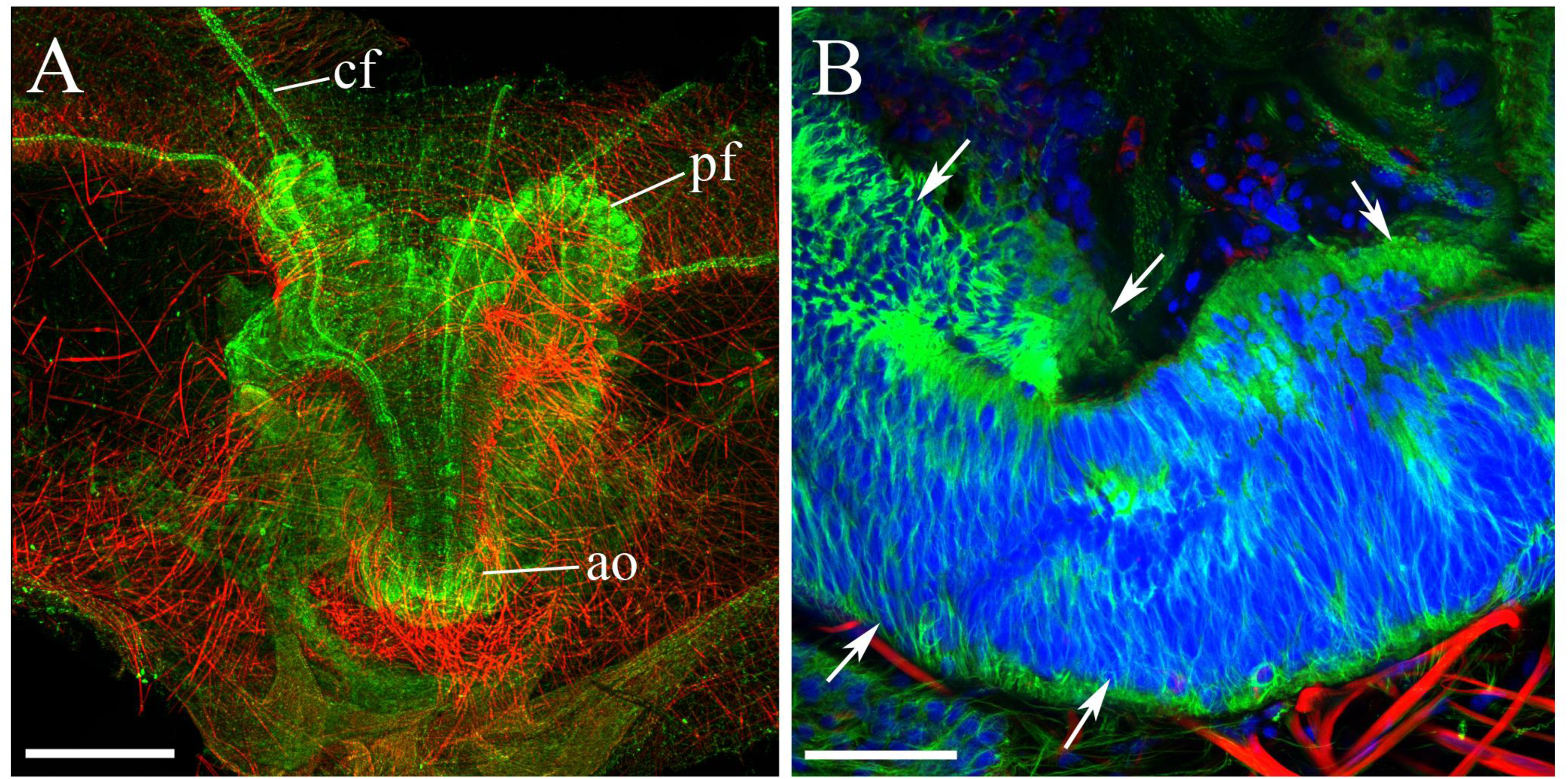
Structure of the aboral organ as revealed by tubulin antibody (green) and phalloidin labeling (red) with nuclear DAPI staining in blue. **A:** Partially retracted aboral complex with the aboral organ and polar fields stained by tubulin antibody; numerous surrounding muscle fibers are shown in red. **B:** Bright tubulin-related fluorescence (green) of the aboral organ is related to many tightly packed small immunoreactive cells (some of them are indicated by arrows). Abbreviations: *ao* - aboral organ; *pf* - polar field; *cf* - ciliated furrow. Scale bars: A - 400 µm; B - 40 µm.

### 3.3 Two Neural Subsystems in *Beroe abyssicola*

Tubulin IR revealed two major parts of the diffused nervous system in *Beroe:* (i) subepithelial polygonal neural network, which covers the entire surface of the body, as well as the internal surface of the entire pharynx, and (ii) a system of diffuse meshwork of neurons and fibers in the mesoglea within the internal gelatinous part of the body.

#### The subepithelial neural network

was similar to the nerve net described in *Pleurobrachia* (Jager et al., 2011; Norekian and Moroz, 2018) and consisted of a distributed polygonal mesh of neural processes, which formed units of different sizes (between 30 and 200 µm) and shapes limiting mostly to quadrilateral (four corners), pentagonal (five corners), hexagonal (six corners), heptagonal (seven corners), and sometimes octagonal (eight corners) configurations (Figs. 6A, 7). Individual strands of the net were composed of many closely packed neuronal processes (Fig. 6C, D).

DAPI staining revealed numerous nuclei inside the network strands suggesting the existence of neuronal somata among neurites (Fig. 7C; Table 1). Consequently, we identified neuronal cell bodies (7-10 µm in diameter) located between the strands of the mesh in the same focal plane (Fig. 6B, C, D). These neurons had one, two, or three processes, which contributed to the structure of the polygonal net (Fig. 6B, C, D; Table 1).

**Figure 6.**
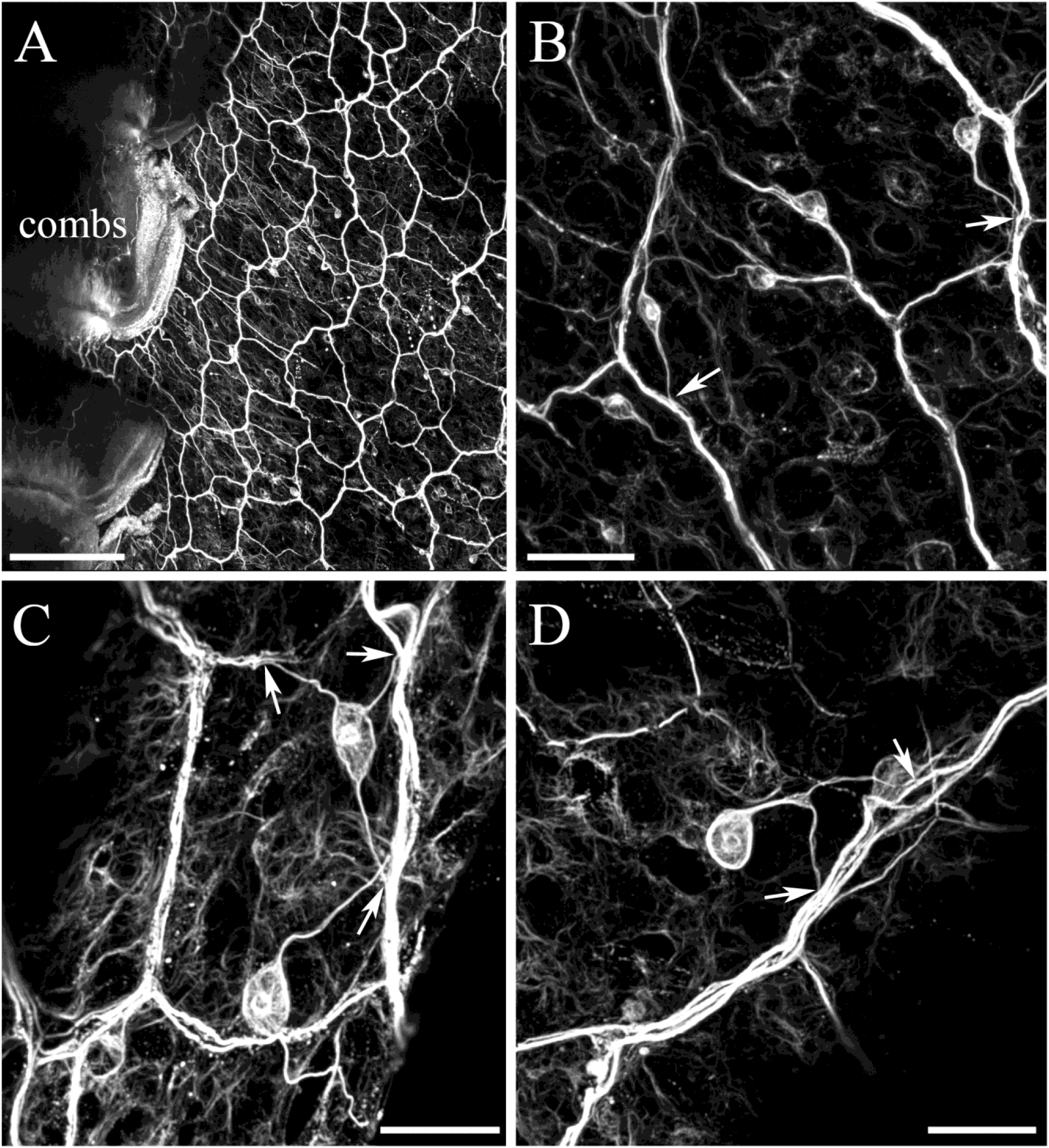
Subepithelial neural network stained with tubulin antibody. **A**: A neural net consists of polygonal units of different shapes and sizes. On the left side, the net connects to the comb row. B, **C,** D: There are individual neurons with one, two or three neurites (arrows) joining the network strands. The strands of the neural net are composed of tightly packed thin processes, rather than a single axon **(D).** Scale bars: A - 200 µm; **B -** 40 µm; C, **D** - 20 µm.

**Figure 7.**
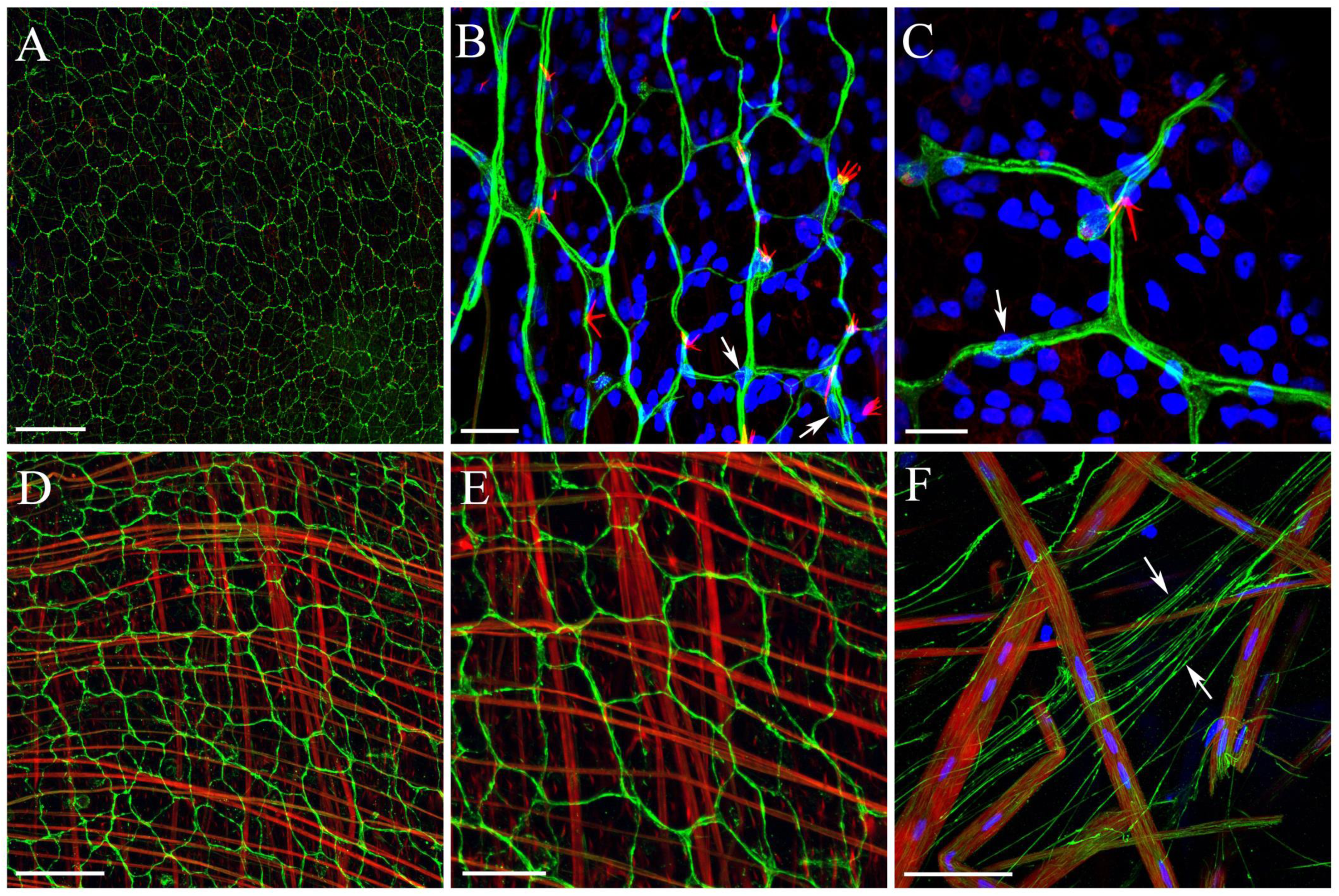
The organization of the subepithelial neural network and adjacent layers (tubulin IR - green; phalloidin –red; nuclear DAPI staining– blue).**A, B, C**:The neural network on the outside surface of the integument consisting of polygonal units of different shapes and sizes and covering the entire surface of the body wall. DAPI reveals neurons inside the strands of the polygon with the central nucleus (arrows). Phalloidin labels some receptor cilia (seetext). **D, E**: Similar organization of the subepithelial network inside the pharynx wall. Phalloidin labels the adjacent large muscle fibers in the mesoglea. **F**: A part of the subepithelial net is a web of fine processes (arrows) running towards the thick muscle fibers and making direct contact with them. Scale bars: A-200µm; B-20µm; C-10µm; D-50µm; E-25µm; F-15µm.

The subepithelial polygonal network covered the whole outer surface of *Beroe* (Fig. 7A, B, C), and the entire internal surface of the pharynx (Fig. 7D, E). We did not recognize any principal differences between the structure of the outer surface network and the internal “pharynx” network. Interestingly, we did not find the pharynx nerve net in *Pleurobrachia* (Norekian and Moroz, 2018).

We also discovered numerous fine neuronal processes associated with the subepithelial network, which were extended to the lower layer of adjacent muscles in mesoglea and making direct contact with them (Fig. 7F). Such an organization suggests that subepithelial network controls the contraction of the large smooth muscles in mesoglea, and therefore, body wall movements.

#### Neurons of Mesoglea

were the second major part of the neural system in *Beroe.* Tubulin antibody labeled a large group of neurons and thin neuronal processes, which spread over the entire mesogleal region (Fig. 8A). All cell bodies had a size varying between 6 and 15 µm depending on the size of the animals. Three types of neurons can be recognized, based on their processes pattern. The first and the most numerous type was represented by bipolar neurons with two processes on opposite sides of the body (Figs. 8B, C; 9A, B). The second type is multipolar neurons with some thin processes projecting in different directions (Figs. 8D; 9C, D). And the third type of cells had long thin filaments cross-crossing the large areas of the mesoglea.

SEM scanning also revealed numerous thin neuronal-like processes among the much thicker muscle fibers in mesoglea of *Beroe* (Fig. 20B, D). These were the same neuronal-like filaments observed during the tubulin antibody staining.

**Figure 8.**
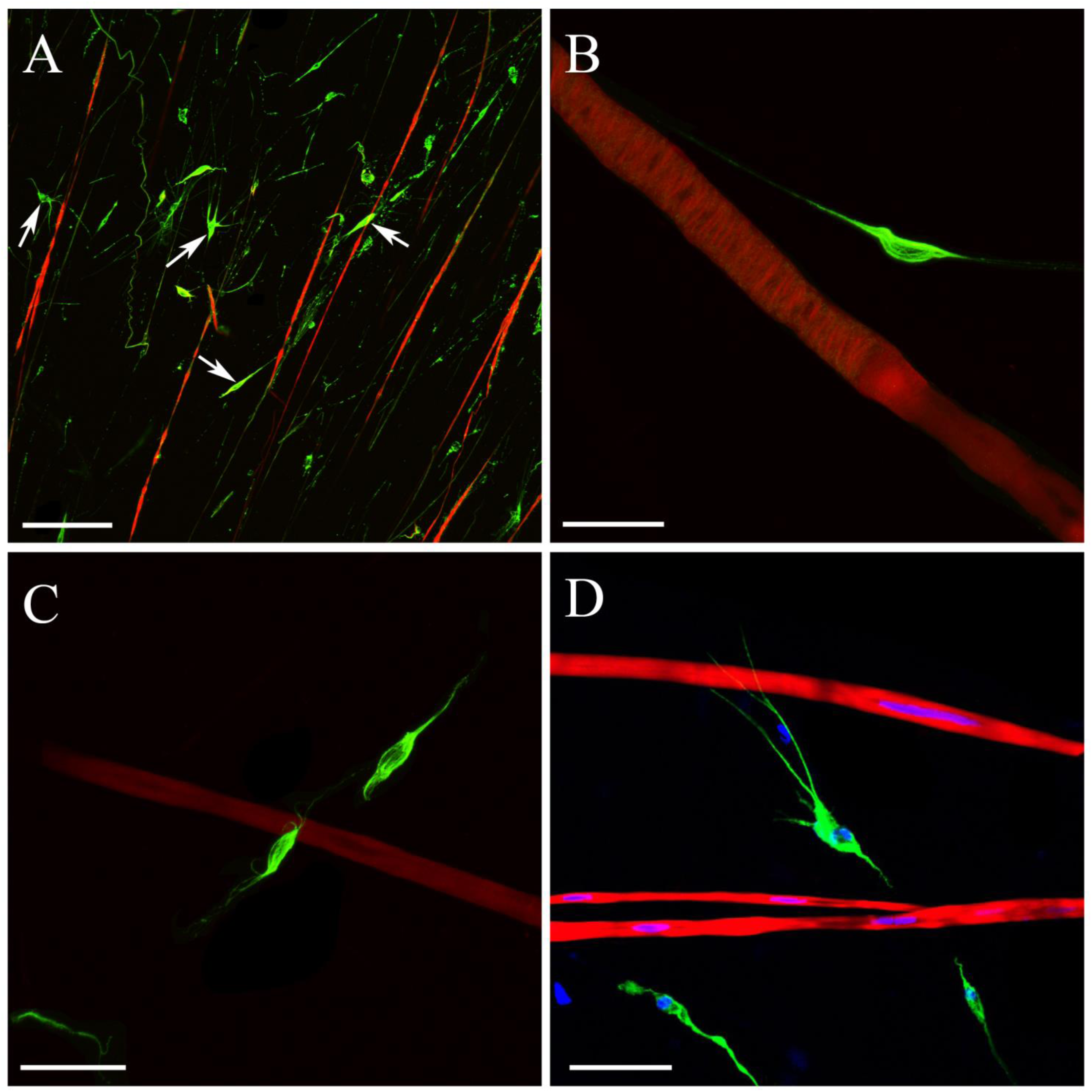
Mesogleal neurons. **A**: The mesoglea area is very rich in muscle and neural cells as well as their processes. Phalloidin (red) stains large muscle fibers, while tubulin antibody (green) labels many small neuronal cell bodies (arrows) as well as thin neuronal-like processes. **B, C:** Neuronal somata can be separated into different types based on the pattern of their processes. The most numerous type is bipolar neurons with processes on opposite poles of the cell. **D:** The second type is represented by multipolar cells with thin processes projecting in different directions. Scale bars: A - 100 fim; B, C - 30 fim; D - 20 fim. D - 20 µm.

## 3.4 Identification of putative receptors in *Beroe*

### Integument/Surface Receptors

We identified five types of putative sensory cells in *Beroe* (Figs. 10, 11, 12, 13 and Table 2). Three types were the surface receptors, which were visualized by both SEM scanning and phalloidin/tubulin IR labeling. The other two types were integrated parts of the subepithelial neural network, which did not open to the surface and could not be seen in SEM scanning.

The receptor type 1 had a tight group of 3 to 9 cilia, 4-8 µm length, projecting in different directions but connected to a single base. They were the most numerous receptors observed during SEM scanning (Fig. 10A). These cilia were brightly stained by phalloidin, while tubulin antibodies usually labeled the cell somata with a large nucleus in the middle (Fig. 11A, B).

**Table 2.** Receptor/sensory cell types identified in *Beroe abyssicola.*

|  | *Receptor type* | *Location* | *Number of cilia* | *Visualization* | *Figures* |
| --- | --- | --- | --- | --- | --- |
| 1. | Receptor with a group of 3-9 cilia | Epithelial; outer surface and pharynx | 3-9 cilia | SEM & Phalloidin | Figs. 10A; 11A, B; 12B; 13C, D |
| 2a. | Receptor with a large thick cilium | Epithelial; outer surface and pharynx | 1 cilium | SEM & Phalloidin | Figs. 10B; 11C; 12B; 13B |
| 2b. | Receptor with a large thick cilium<br>Subtype in the "lips" area | Epithelial; lips and outside mouth | 1 cilium | SEM & Phalloidin | Figs. 14, 15 |
| 3. | Receptor with a long thin cilium | Epithelial; outer surface and lips | 1 cilium | SEM | Figs. 10C; 14C |
| 4. | Ciliated neurons between strands of the subepithelial net | Subepithelial; a part of the neural net | 1 cilium | Tubulin IR | Figs. 11D, E |
| 5. | Receptors with tubulin IR labeled processes joining the subepithelial net | Subepithelial; a part of the neural net | no cilia | Tubulin IR, only processes labeled | Figs. 11D, E |

**Figure 9.**
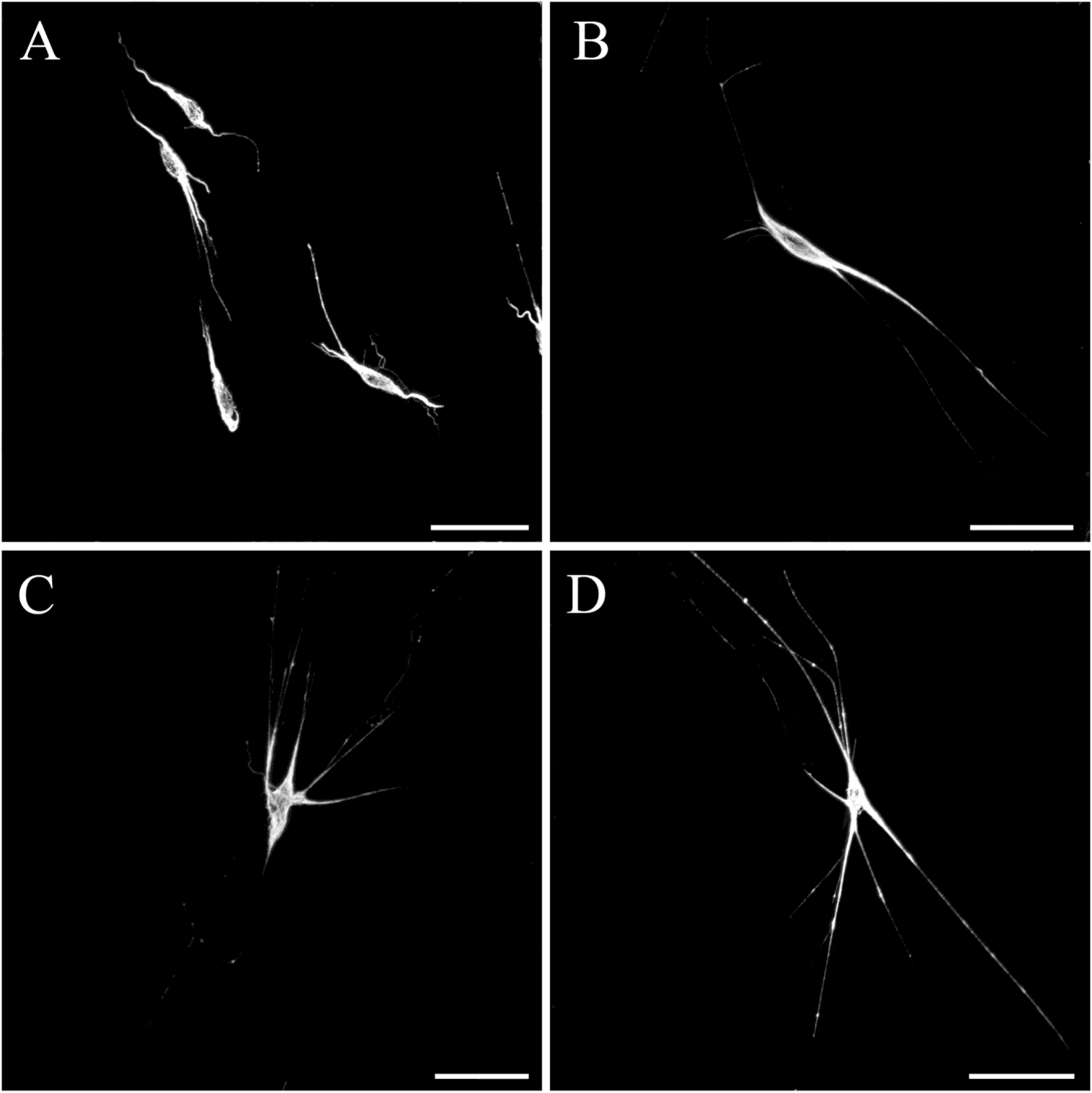
Mesogleal neurons stained by tubulin antibody. **A, B:** Bipolar neural cells. **C, D:** Multipolar neurons. Scale bars: A, B, C - 30 pm; D - 40 µm.

**Figure 10.**
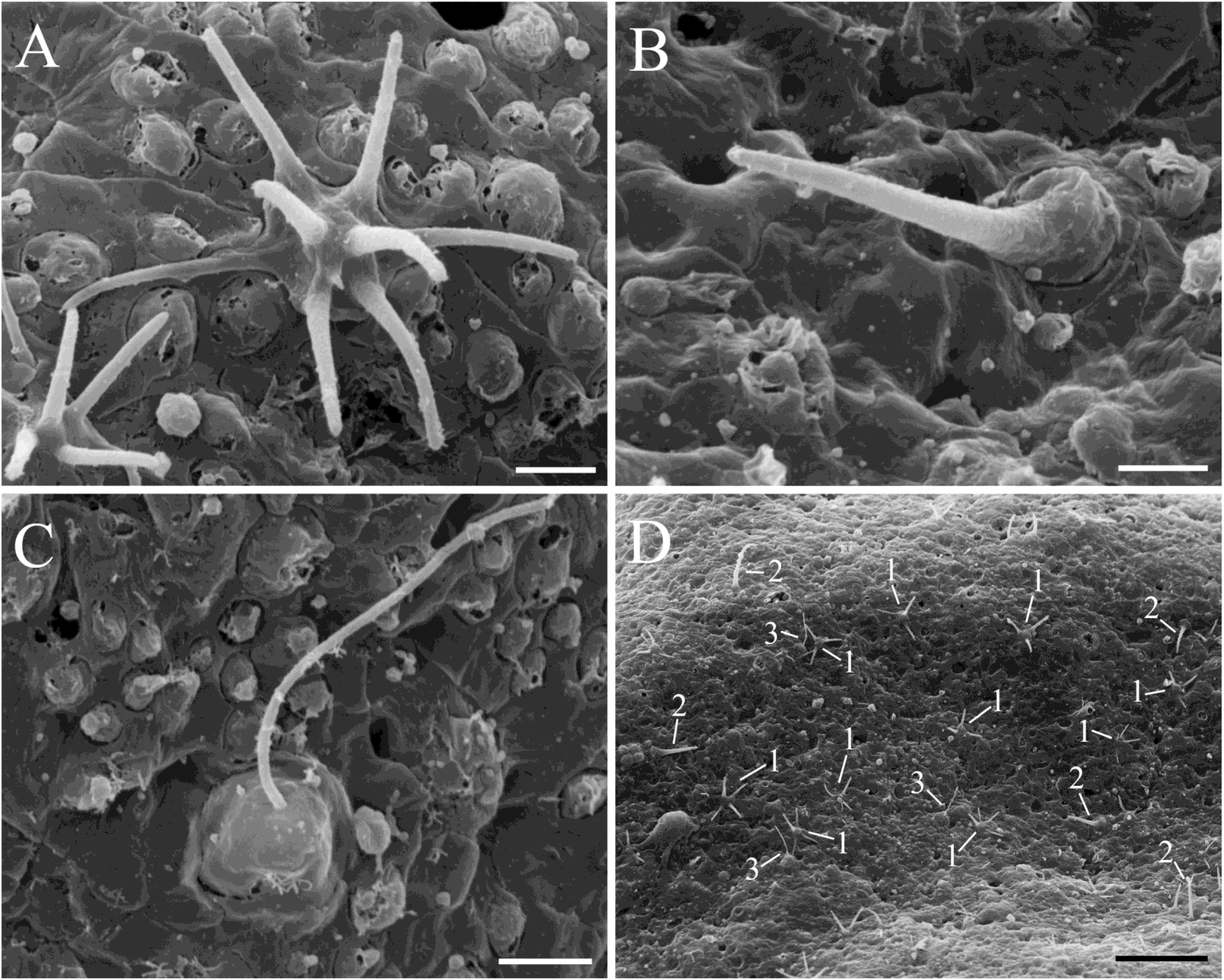
SEM of different ciliated receptors on the surface of the body. **A**: Receptors with a group of 3 to 9 cilia on the top of a single round base. **B:** Receptor with a single large and thick cilium. **C:** Receptor with a single thin cilium on the apical part of a cell. **D:** Distribution of all three receptor types on the skin surface of *Beroe.* The most numerous is the receptor type 1 with multiple cilia. Fewer density of the receptor type 2 (with a long thick cilium) and the receptor type 3 (thin cilium). Scale bars: A, B, C - 2 µm; D-20µm;.

**Figure 11.**
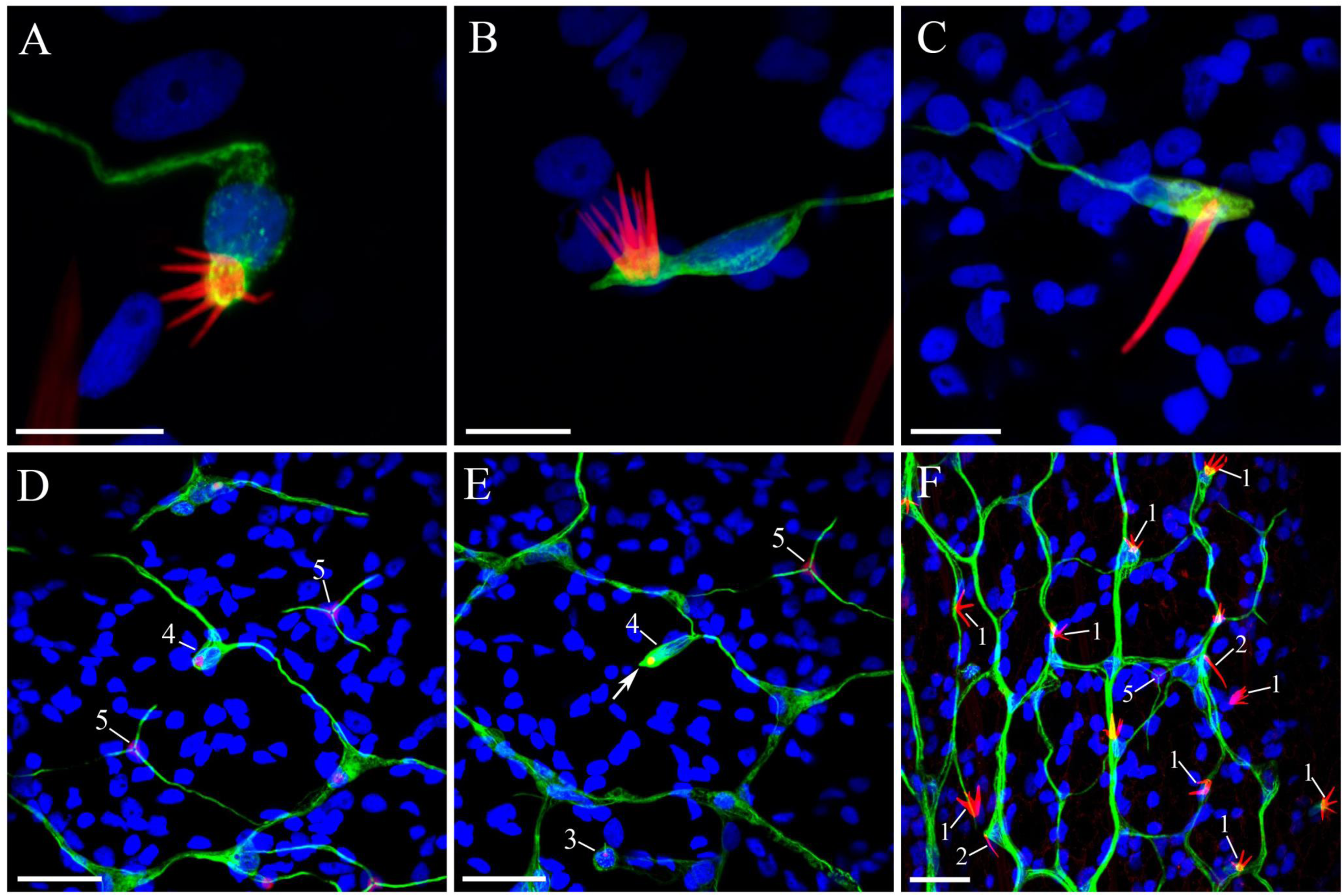
Five receptors types are associated with the subepithelial neural net on the surface of the body. (green - tubulin IR; red - phalloidin; blue - nuclear DAPI labeling). **A, B**: The first type of receptors contains a compact group of 3 to 9 long cilia labeled by phalloidin. The cell body of the receptor is labeled by tubulin antibody and projects a single processes. **C**: The second receptor type has a large and thick cilium labeled by phalloidin on the top of a tubulin IR cell with a neurite. **D, E**: The third type of receptors is difficult to see since it has a very thin cilium that is weakly stained by tubulin antibody only (3). The fourth receptor type includes sensory cells with a short cilium stained with tubulin antibody (arrow); these receptors have two or three processes within the neural net similarly to other neurons in the net (4). The fifth type of receptor cells has three radial tubulin IR neurites fusing with the subepithelial neural net, but with the cell body of the receptor was not labeled by the tubulin antibodies (5). **F**: Distribution of different receptor types on the outer surface. Scale bars: A, B, C - 10 μm, D, E, F - 20 μm.

**Figure 12.**
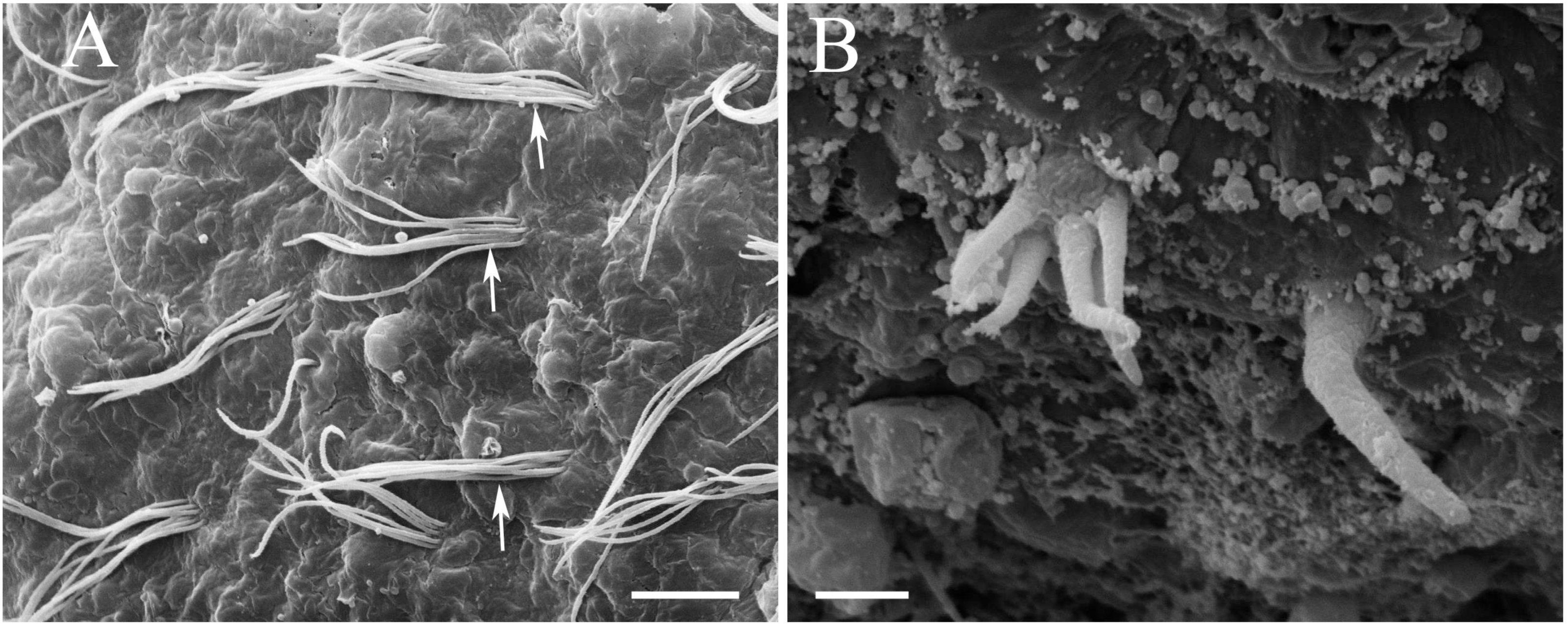
SEM of the pharynx surface. **A**: Groups of thin cilia (arrows) outlining the inner surface of the pharynx. **B:** Two types of receptors were identified - multiciliated receptors and receptors with a large and thick cilium. Scale bars: A - 5 µm; B - 2 µm.

**Figure 13.**
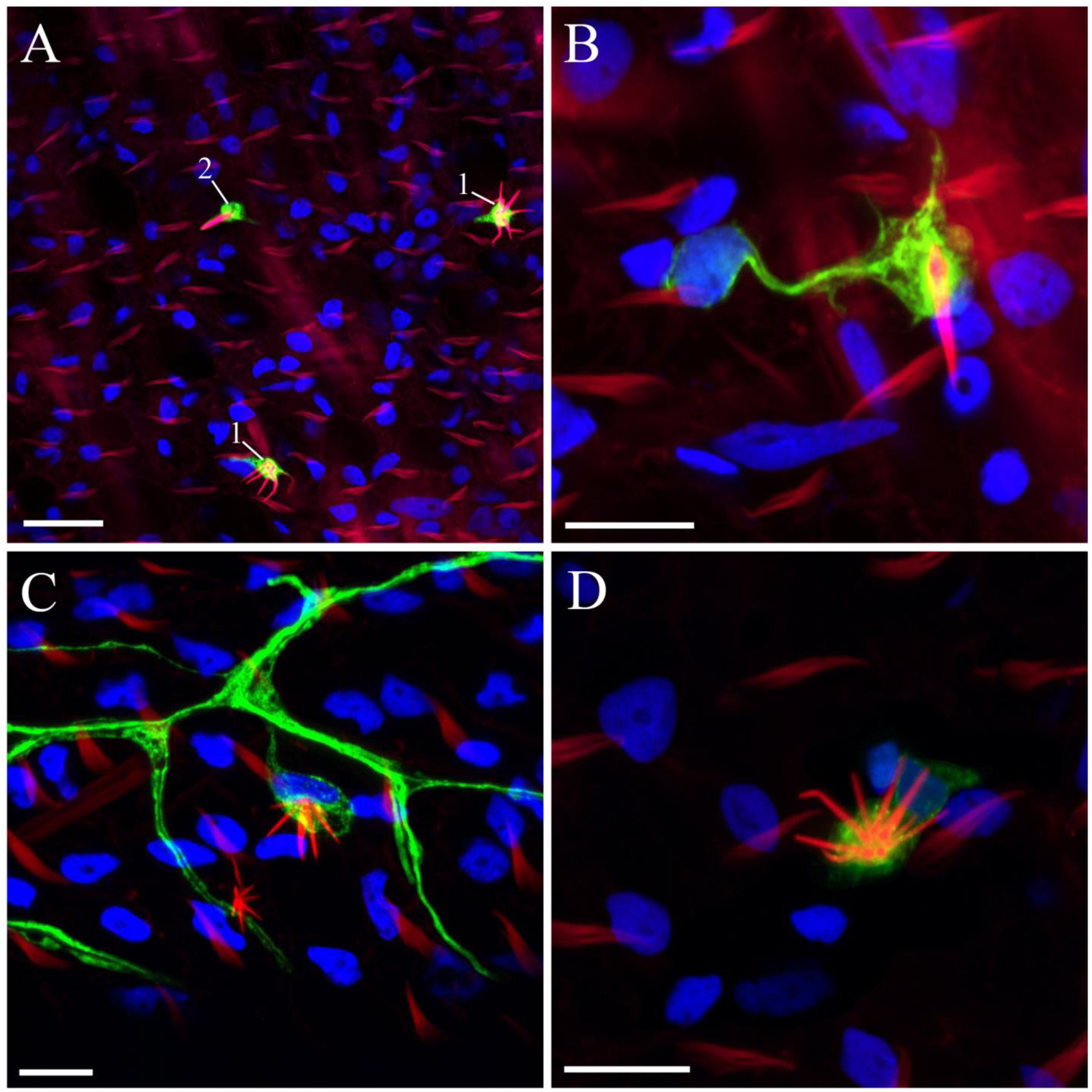
Two types of receptors are associated with the subepithelial neural net on the inner surface of the pharynx. (green - tubulin IR; red - phalloidin; blue - nuclear DAPI labeling). **A:** The density of the receptors inside the pharynx is several times lower than on the outer surface. **B:** Receptors with a single large cilium labeled by phalloidin. **C, D:** Another type of receptors with a group of 3 to 9 cilia labeled by phalloidin. Scale bars: A - 20 µm; B, C, D - 10 µm.

The receptor type 2 had a long and thick single cilium (up to 9-10 µm length) at a wide base (Fig. 10B). This cilium was brightly stained by phalloidin, while tubulin IR was detected in the cell body (Fig. 11C). These receptors were less numerous than the first type.

The third receptor type had a single, very thin cilium, up to 8 µm length, located at the apical part of about 5 µm cells (Fig. 10C). We identified this receptor type mostly in SEM imaging. Its cilium was weakly marked by tubulin IR (Fig. 11E) - a similar receptor type was also found in *Pleurobrachia* (Norekian and Moroz, 2018). This receptor type was less numerous than the type one (Fig. 10D). In general, the density of all types of receptors was higher near to the aboral organ and polar fields compare to the skin area between the comb rows.

The fourth receptor type was represented by neuron-like sensory cells located between the strands of the subepithelial neural network (Fig. 11D, E). These cells were almost indistinguishable from other neurons within the polygonal mesh described earlier (Fig. 6B, C, D). They had the cell bodies about 7-10 µm in diameter and one, two, or three processes joining the strands of the subepithelial network. The main difference was a very short cilium on the top of their cell body brightly stained by tubulin IR (Fig. 11E); the base of the cilium was always labeled by phalloidin (Fig. 11D).

The fifth type of receptors included cells located slightly below the plane of the main network. These cells usually had three radial tubulin IR neurites fusing with the subepithelial network (Fig. 9D, E). However, the cell body itself was not tubulin IR. A large nucleus and few short phalloidin-stained pegs were always observed in the point of conversion of three neurites (Fig. 9D, E).

### Pharynx receptors

Receptors were found not only on the outer surface of the body but also throughout the pharynx. The entire surface of the pharynx was covered with groups of long thin cilia, which might support the movement of ingested food (Fig. 12A). Among those cilia, we have identified two types of receptors: (i) the multiciliated cells, and (ii) the cells with a long thick nonmotile cilium (Fig. 12B, Table 2). The cilia in both types of receptors were stained by phalloidin, while their cell bodies were tubulin IR (Fig. 13). The density of the receptors in the pharynx was by 3-5 times less compared to the body surface.

### Lips/Mouth receptors

Being an active predator, *Beroe* ought to have a significant sensory representation in the oral area (Fig. 14A), which can detect mechanical stimulation and chemical inputs from preys. The very edge of the mouth in *Beroe* contains a clearly defined narrow band (Fig. 14A), which can be morphologically identified as “lips”. Here, we identified two types of receptors in the “lips” area.

A group of cells, also known as sensory “bristles” (Tamm and Tamm, 1991a), was similar to the receptor type 2 on the body surface (Figs. 10B, 11C; Table 2). These receptors were easily identifiable because of their very long thick cilium with narrowing end. The densest concentration of these receptors was found on the outside edge of the “lips” (Fig. 14B, C), but could be seen throughout the entire “lips” area among other cilia and papillae, although at lower density (Fig. 14D). The cilium of the type 2 receptor was brightly stained by phalloidin (Fig. 15A, B, C), and could be observed inside the mouth among the feeding macrocilia (Fig. 15A). Just outside the mouth, we identified many small sensory bulges (or bumps), which were densely covered with sensory cilia of the type 2 receptors (Fig. 14A, 15A, B). These areas might provide important sensory information when “lips” are closed during *Beroe” s* search for food. The type 3 receptor, with a long single cilium, was also found at high density on the outer edge of the “lips” (Fig. 14C) and were identified only via SEM scanning.

**Figure 14.**
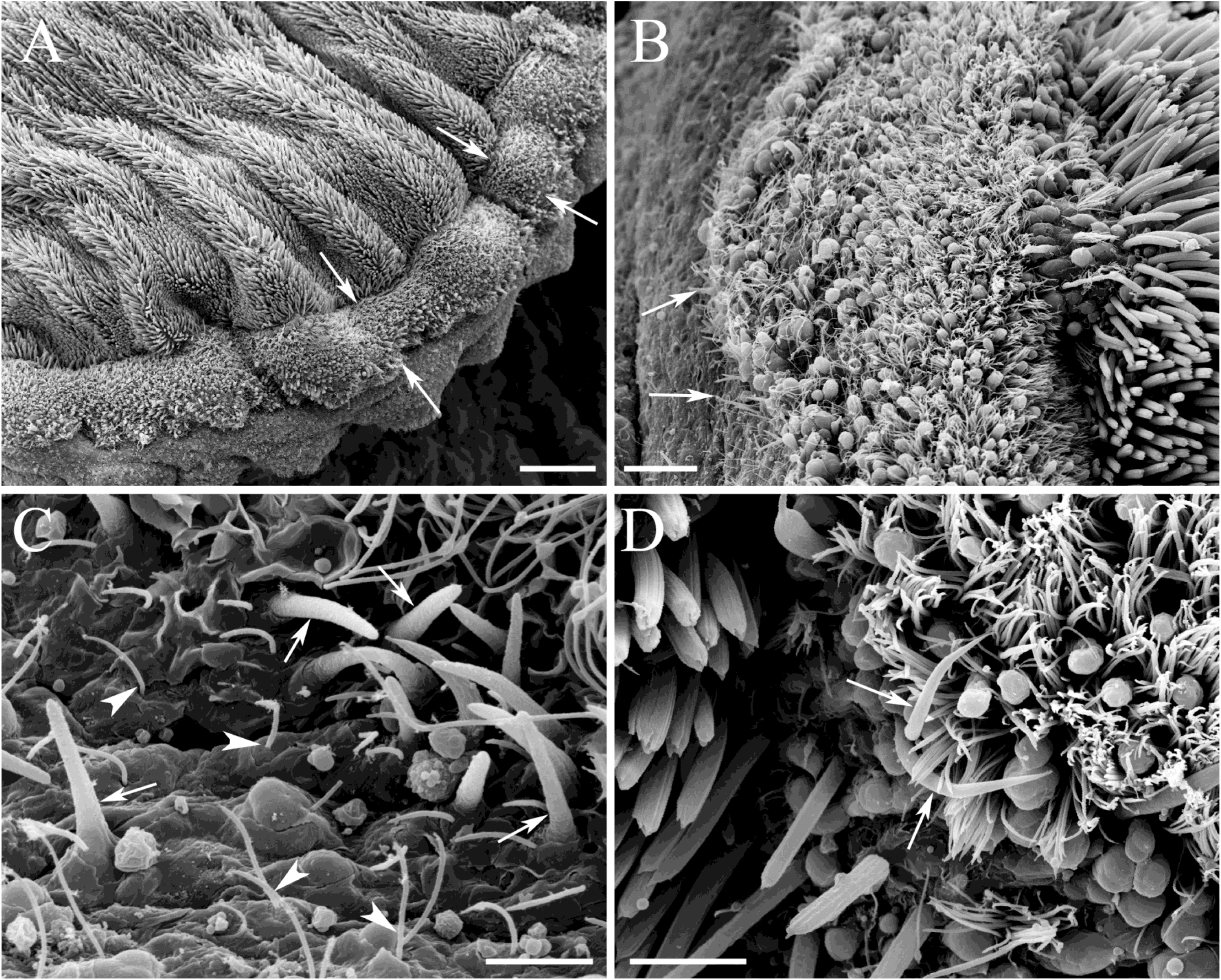
SEM of ciliated receptors on the “lips.” **A:** The edge of the mouth in *Beroe* has a well-defined narrow band - the “lips” (arrows). **B, C:** The “lips” contain many receptors with a single large and thick cilium. The highest concentration of those receptors was found on the outside edge of the “lips” (arrows). Numerous receptors with a thin cilium are also present in the same area (arrowheads). **C:** Receptors with a long thick cilium are easily identifiable throughout the “lips” area including the side facing the pharynx (arrows). Scale bars: A - 100 μm; B - 20 μm; C - 5 μm; D - 10 μm.

**Figure 15.**
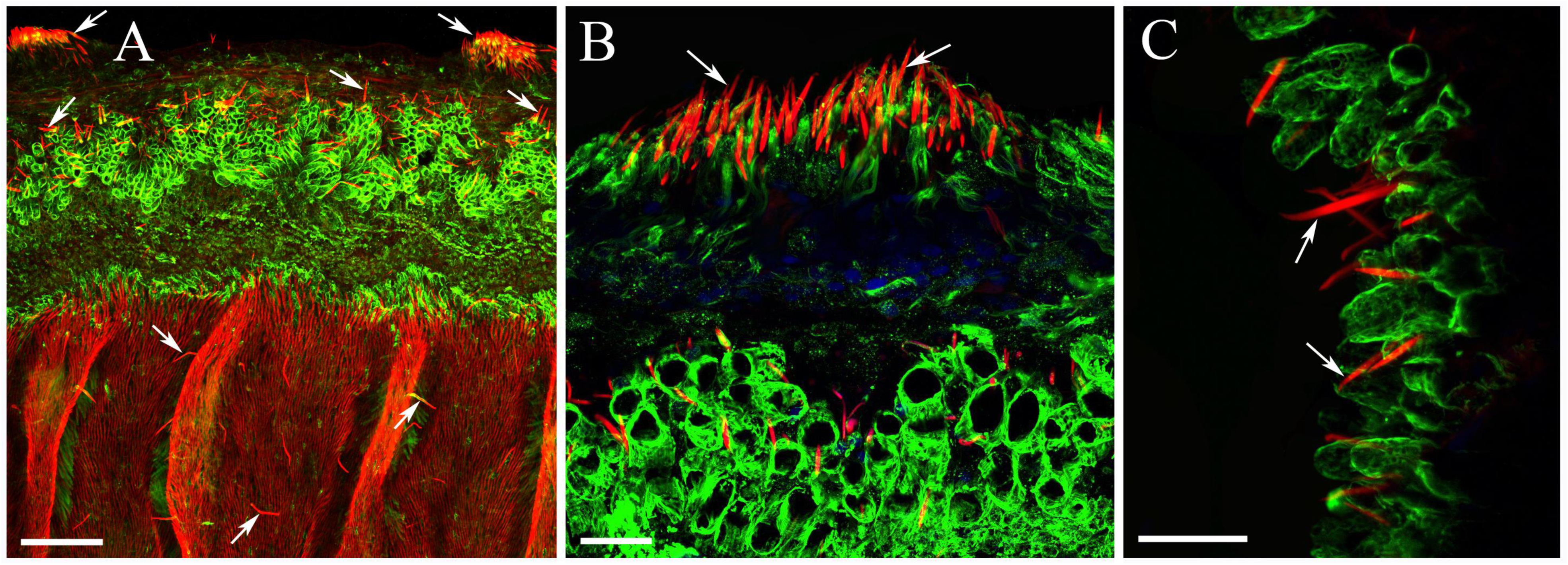
Phalloidin-labeled (red) sensory cilia in the mouth area. **A**: Numerous receptors with a single large cilium brightly stained by phalloidin (arrows) are found on the “lips,” inside the mouth and in large congregates on the outside surface. **B:** Sensory bulges densely covered with phalloidin labeled cilia (arrows) on the body surface outside the “lips.” **C:** Sensory cilia on the outer edge of the “lips.” Scale bars: A - 100 µm; B, C - 20 µm.

#### 3.5 Ciliated Structures of Mouth and Pharynx

The life of ctenophores is primarily based on cilia, often with unique lineage-specific adaptations (Tamm, 2014). *Beroe,* as a planktonic macro-feeder, preys on large soft-bodied ctenophores. The distinct characteristic of Beroida is a set of specialized feeding-related cilia on the inner side of the mouth called macrocilia (Horridge, 1965c; Tamm, 1982; Tamm, 1983; Tamm and Tamm, 1984; 1985; 1987; Tamm and Tamm, 1991a; b). The macrocilia are sharp and stiff enough to tear down the soft tissue of prey and function like teeth. Each macrocilium consists of an array of individual ciliary axonemes, which rootlets penetrate and anchor the massive bundle of non-contractile actin filaments extending throughout the length of the cell suggesting that these actin filament bundles “may serve as intracellular tendons” (Tamm and Tamm, 1987; Tamm, 2014).

Using SEM imaging, we found that the density of macrocilia was the highest closer to the outer edges of the mouth and then slowly decreased toward the pharynx and eventually disappeared (Fig. 16C, D). The macrocilia were about 10-20 µm long and 2-4 µm in diameter (Fig. 16E F). At the outer edges of the mouth, the macrocilia were always straight usually displaying three sharp teeth (Fig. 16F). Further into the pharynx, the macrocilia had only one sharp end and were bent like a hook pointing towards the inside, and presumably forcing the food to move only in one direction (Fig. 16E).

The macrocilia itself were tubulin IR (Fig. 17C, D), and always attached to the phalloidin-labeled bundles of actin filaments (Fig. 17C, D). Those bundles of actin filaments extended 5-10 µm and had a shape of a stretched teardrop.

**Figure 16.**
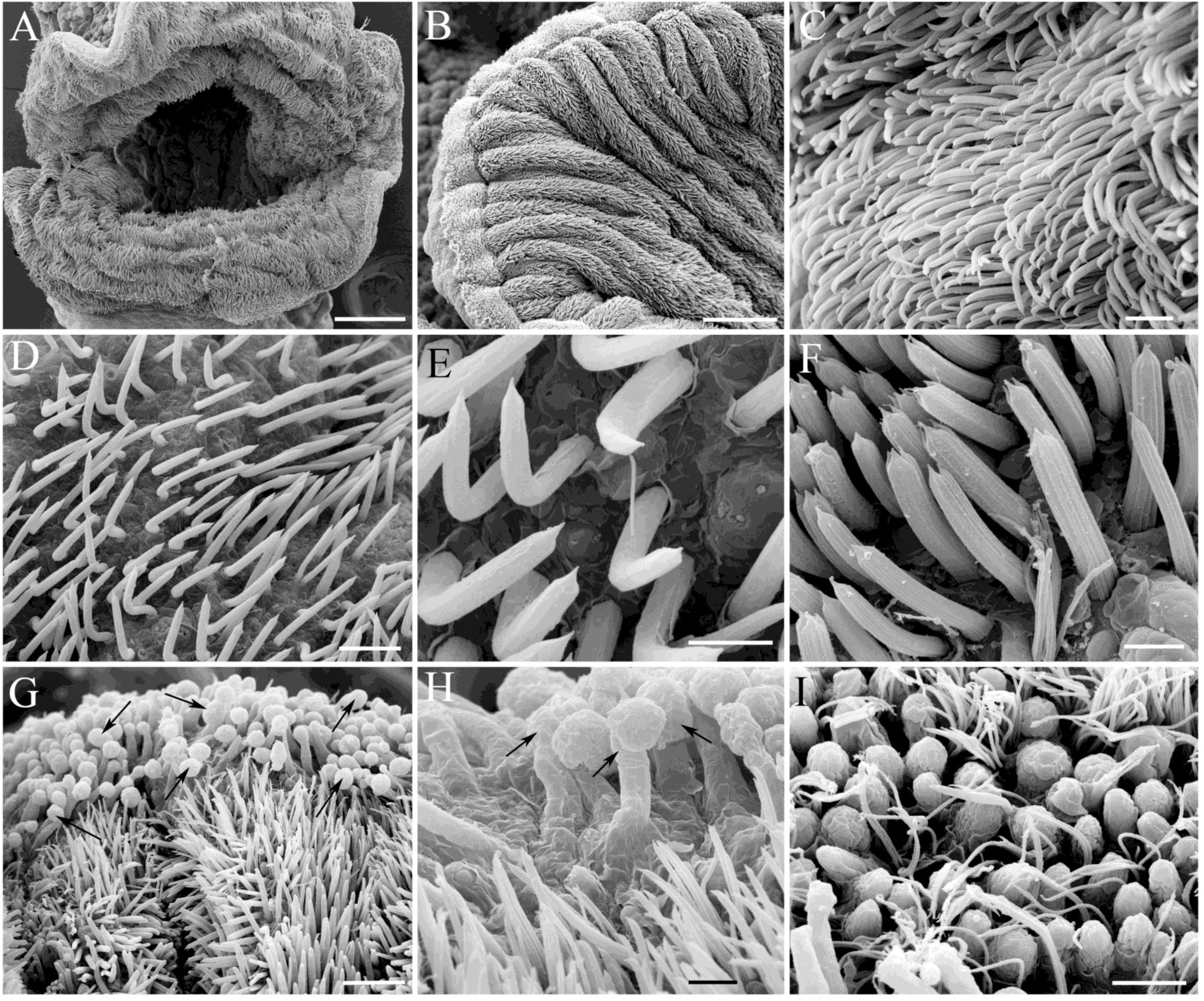
SEM of the mouth area. **A**: Wide elongated mouth. **B:** The entire area inside the mouth up to the “lips” is covered with macrocilia. **C:** Macrocilia density is higher closer to the outer edge of the mouth. **D:** Further into the pharynx the macrocilia density is reduced. **E:** Deep into the pharynx, the macrocilia have a single sharp end, and they usually bent like a hook pointing towards the inside of the pharynx. **F:** Closer to the outer edges of the mouth, the macrocilia are always straight and display three sharp teeth at the top. **G, H:** Papillae-like structures (arrows) in the “lips” area, which are the part of the “lips” reversible adhesion mechanism in *Beroe* (see the text). **I:** Some of the “locking” papillae are shorter than others and are spread over the entire “lips” area. Scale bars: A, B - 200 µm; C, D - 10 µm; E, F - 5 µm; G - 20 µm; H, I - 5 µm.

**Figure 17.**
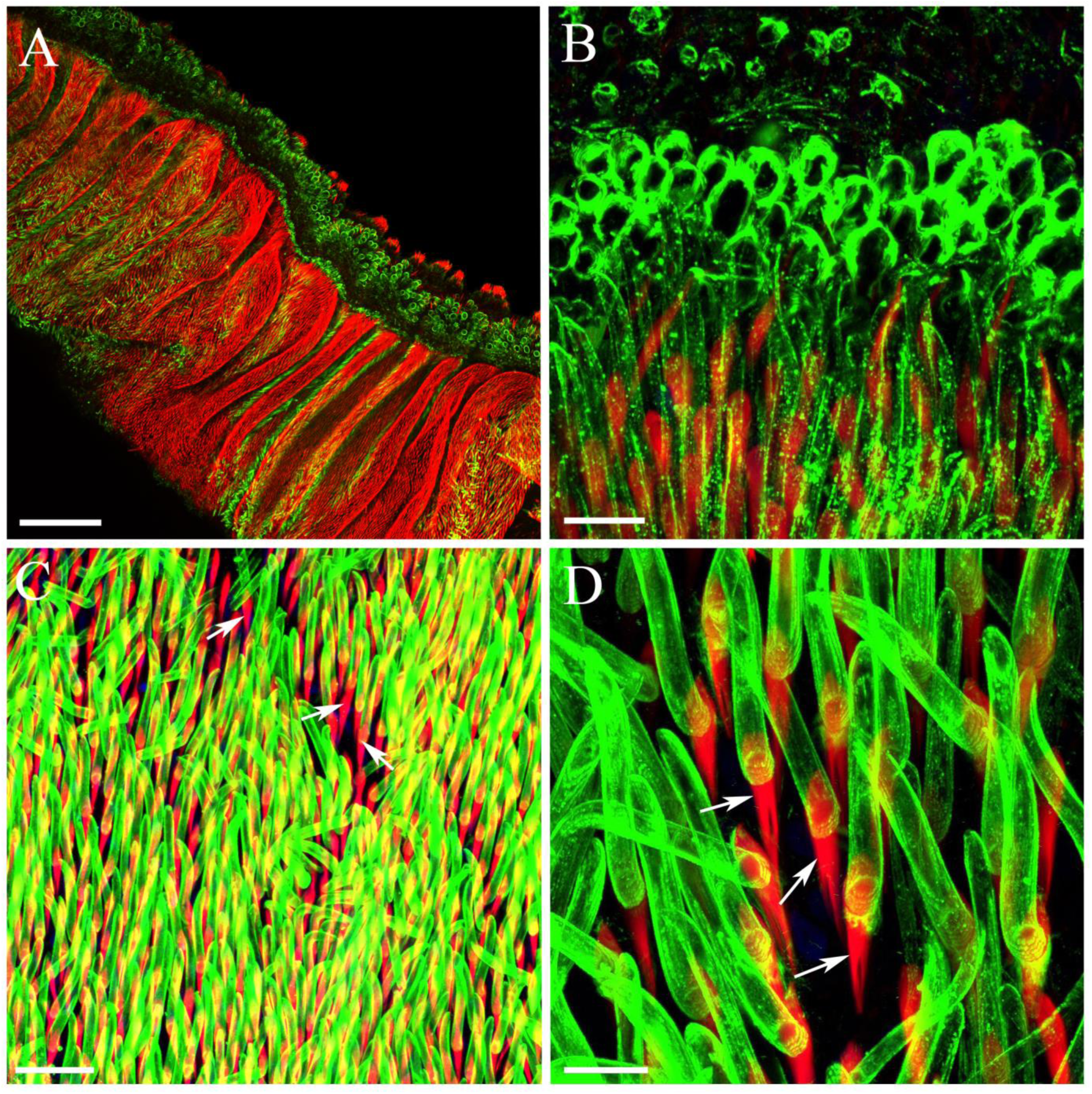
Phalloidin (red) and tubulin antibody (green) labeling in the mouth area. **A, B:** The “locking” papillae in the “lips” are labeled by tubulin antibody. **C, D:** The long cylindrical body of macrocilia itself are stained by tubulin antibody and attached to the phalloidin-labeled bundles of actin filaments (arrows) inside the macrociliary cell. Scale bars: A - 300 µm; B, D - 10 µm; C - 30 µm.

Unless they are feeding, *Beroe* keep their mouth closed by paired strips of adhesive epithelial cells located on the opposing lips (Tamm and Tamm, 1991b; a; Tamm and Tamm, 1991c; Tamm, 2014). The interlocking of the cell surfaces is thought to mechanically fasten the paired adhesive strips together like a jigsaw puzzle. Contact with prey triggers a muscular separation of the lips.

We have identified numerous papillae-like structures, which were the part of that “lips” locking mechanism in *Beroe* (Fig. 16G, H, I). They could reach up to 10-15 µm height with the round-shaped head of about 5 µm in diameter. Some of them were shorter with only round-shaped heads elevating above the epithelium. These papillae were spread over the entire “lips” area among the sensory cilia (described above) and groups of thin and motile cilia presumably responsible for the flow of water or maybe even having some sensory function (Fig. 16I). These locking papillae were tubulin IR (Fig. 17A, B) as many epithelial secretory cells in *Pleurobrachia* (Norekian and Moroz, 2018).

#### 3.6 Meridional canals and Ciliated Rosettes

The pharynx in *Beroe* leads to a small stomach (infundibulum) located close to the aboral organ, and then into the eight meridional canals running under the comb rows and a pair of pharyngeal (=paragastric) canals, which extend along the length of the pharynx to the mouth opening. The meridional and pharyngeal canals give off numerous side branches throughout the mesoglea forming a wide gastrovascular system.

The walls of gastrovascular canals and all their branches on the side facing the mesoglea contained numerous ciliated pores called ciliated rosettes (Hernandez-Nicaise, 1991), which were stained by tubulin antibody (Fig. 18).

**Figure 18.**
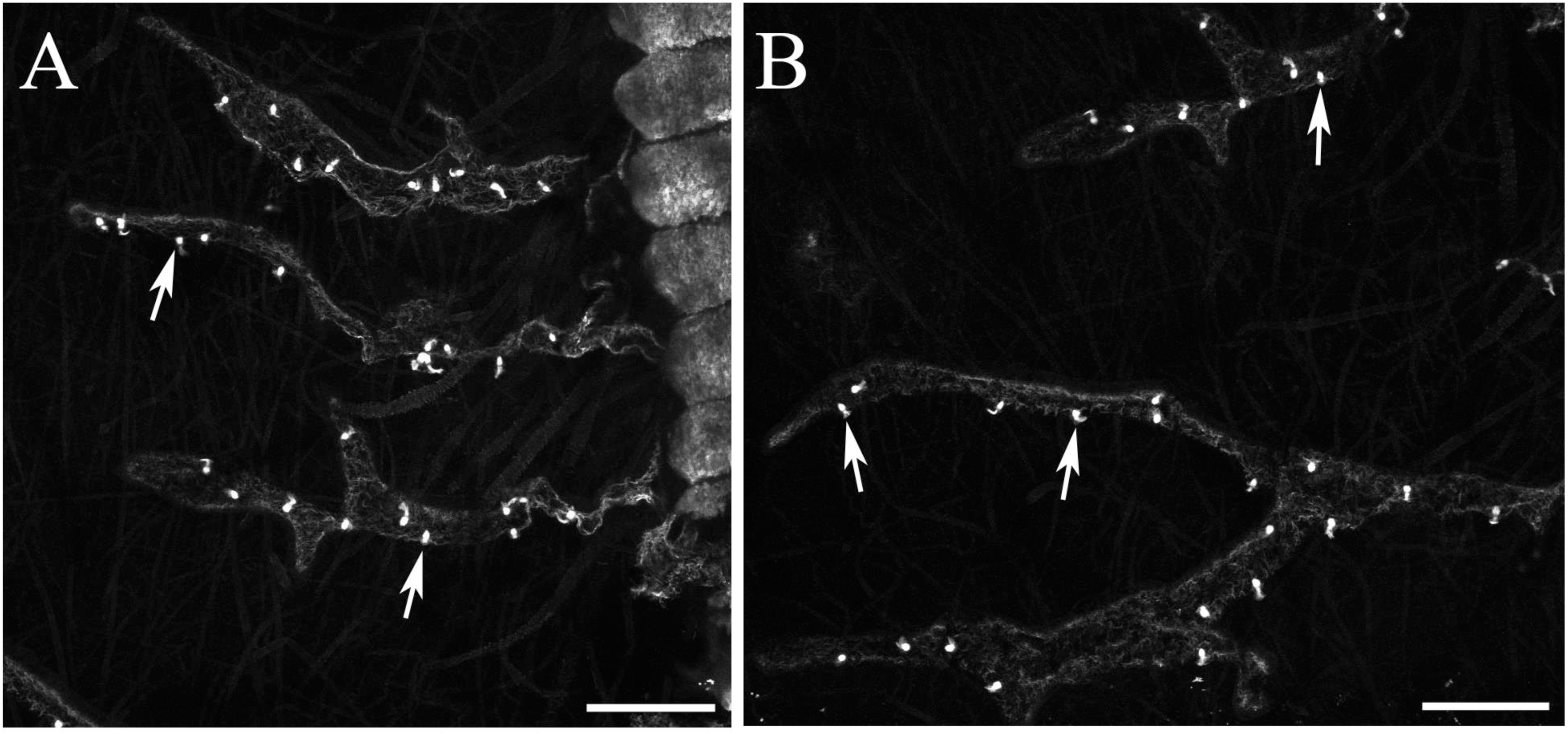
Tubulin IR in the meridional canals of *Beroe.* **A, B**: The branches of meridional canals contain numerous pores called ciliated rosettes, which are brightly stained by tubulin antibody (arrows). Scale bars: A, B - 200 µm.

Each rosette consists of two superimposed rings of eight ciliated cells. The cilia of the upper ring beat into the canal, while the inner ring and its cilia protrude into the mesoglea. Both groups of cilia were tubulin IR (Fig. 19), but the cell bodies of the upper ring were not labeled (only the DAPI-stained nuclei could localize them). The intravascular cilia were short (about 3-4 µm length) and grouped into the conical cluster (Fig. 19B, C, D). On the contrary, cell bodies of the inner ring were stained with tubulin antibody together with their long (up to 15 µm in length) cilia (Fig. 19B, C, D).

**Figure 19.**
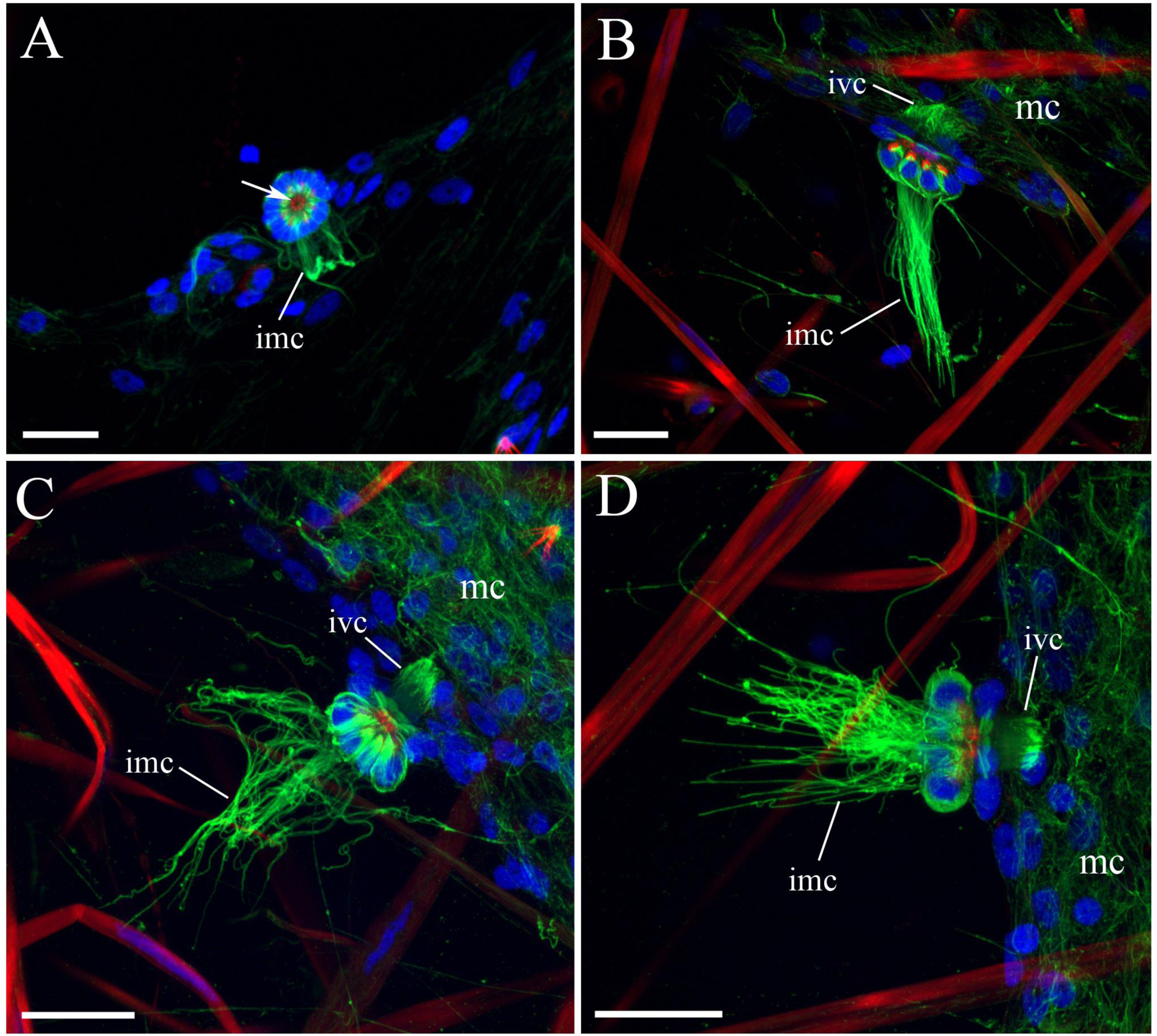
Ciliated rosettes in *Beroe*. (tubulin IR - green; phalloidin - red, DAPI stained nuclei - blue). Each rosette consists of two superimposed rings with eight ciliated cells. Two groups of cilia are labeled by tubulin antibody - shorter intravascular cilia, wnich beat into the meridional canal, while the much longer intramesogleal cilia protrude into the mesoglea. Phalloidin labels the round extensions of the upper ring cells, which form a fibrous ring or diaphragm in the center of a rosette (arrow). Abbreviations: *me -* meridional canal; *imc* - intramesogleal cilia; *ivc -* intravascular cilia. Scale bars - 20 µm.

**Figure 20.**
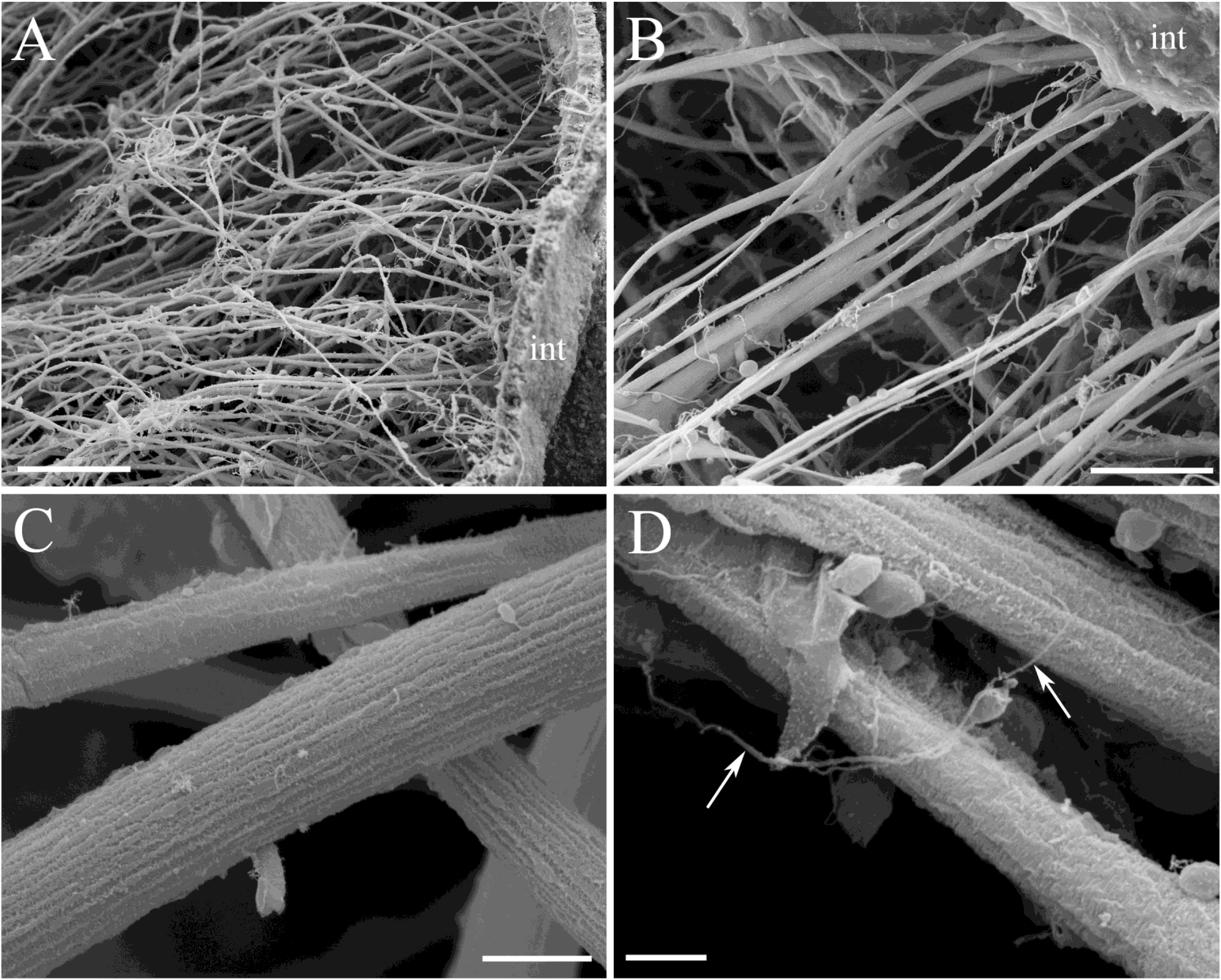
SEM of the muscles in *Beroe*. **A, B**: Mesoglea contains numerous smooth muscle fibers, many of which are firmly attached to the outside integument (int). C: The muscle fibers have a variable thickness. D: In addition to thick muscle fibers, there are many thin neuronal-like processes (arrows). Scale bars: A - 100 µm; B - 50 µm; C - 5 µm; D - 2 µm.

Phalloidin labeled the eight round extensions of the upper ring cells, which formed a fibrous ring or diaphragm in the center of a rosette (Fig. 19). These extensions can retract or extend, therefore, opening or closing the central orifice. The ciliated rosettes thus allow the exchange of fluids between gastrovascular canals and mesogleal compartment with fluids being propelled by the beating of the cilia.

#### 3.7 The diversity of muscle systems in *Beroe*

The mesoglea in *Beroe* contains many giant smooth muscle fibers at much higher density compared to less active *Pleurobrachia* (Norekian and Moroz, 2018). During SEM imaging, we purposefully broke the *Beroe* body wall to expose numerous muscles attached to the outside integument (Fig. 20A). The muscle fibers themselves had a variable thickness between 1 and 8 µm in diameter, which was maintained throughout their entire length (Fig. 20B, C, D). Rarely, we saw bifurcations of single muscle fiber into two branches. There were also numerous thin neuronal-like processes frequently attached to large muscle fibers (Fig. 20D).

Phalloidin staining revealed three groups of muscle cells in the mesoglea: (i) the circular muscles encircling the pharynx and the aboral organ (Fig. 21A, B, C); (ii) the longitudinal muscle fibers (Fig. 21B, C); (iii) shorter radial muscle fibers, which crossed the mesoglea with their branched endings attached to the outside integument, on one side, and the pharyngeal wall on the other (Fig. 21C, D). All muscle cells had multiple nuclei spread along their entire length (Fig. 21D, E).

**Figure 21.**
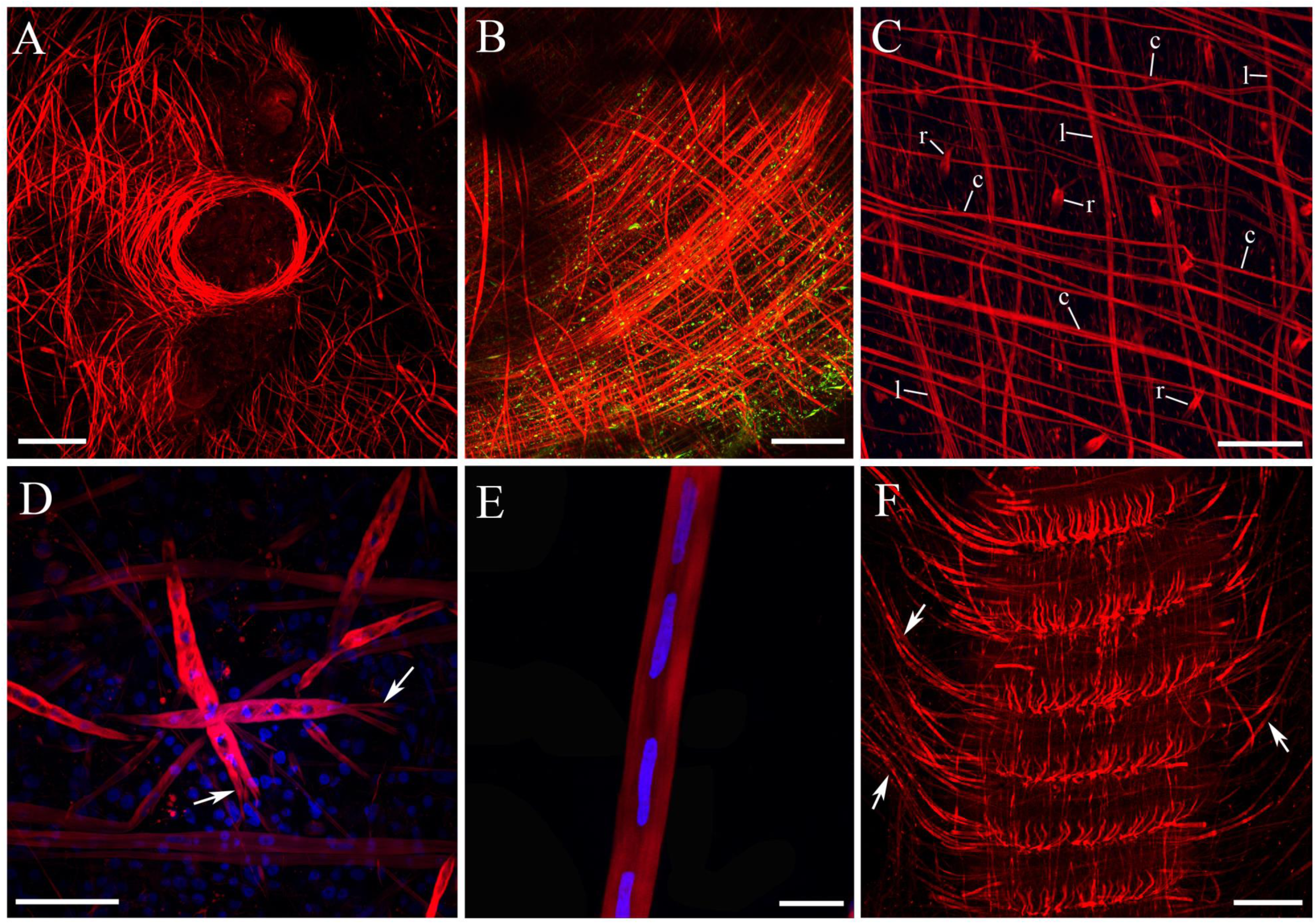
The diversity of muscle groups in *Beroe*. stained by phalloidin (red). **A:** Circular muscles around the aboral organ. **B:** Smooth circular muscle fibers in the mesoglea encircling the pharynx. **C:** The circular *(c)* muscles and longitudinal *(I)* muscles form a loose rectangular mesh of fibers in mesoglea. Radial *(r)* muscles cross the mesoglea and connect the outside integument and pharyngeal wall. **D:** Much shorter radial muscle fibers with multiple nuclei have the branched endings (arrow) at the point of attachment. **E:** Long smooth longitudinal muscle fiber contains multiple nuclei. **F:** Comb rows have the array of very long individual muscles (arrows) attached to them along their entire length and a group of short muscle-like non-contractile elements that connect each comb plate. Scale bars: A - 200 µm, B - 150 µm, C - 100 µm, D - 50 µm; E - 10 µm; F - 200 µm.

Phalloidin staining also revealed large muscles attached to the comb rows, which were evenly spread along the entire row length and penetrated the mesoglea on the other end (Fig. 21F). There were also short non-contractile fibers attached to each comb base, similar to those described in *Pleurobrachia* (Norekian and Moroz, 2018). Table 4 summarizes the information about muscle types in *Beroe*.

**Table 3.**
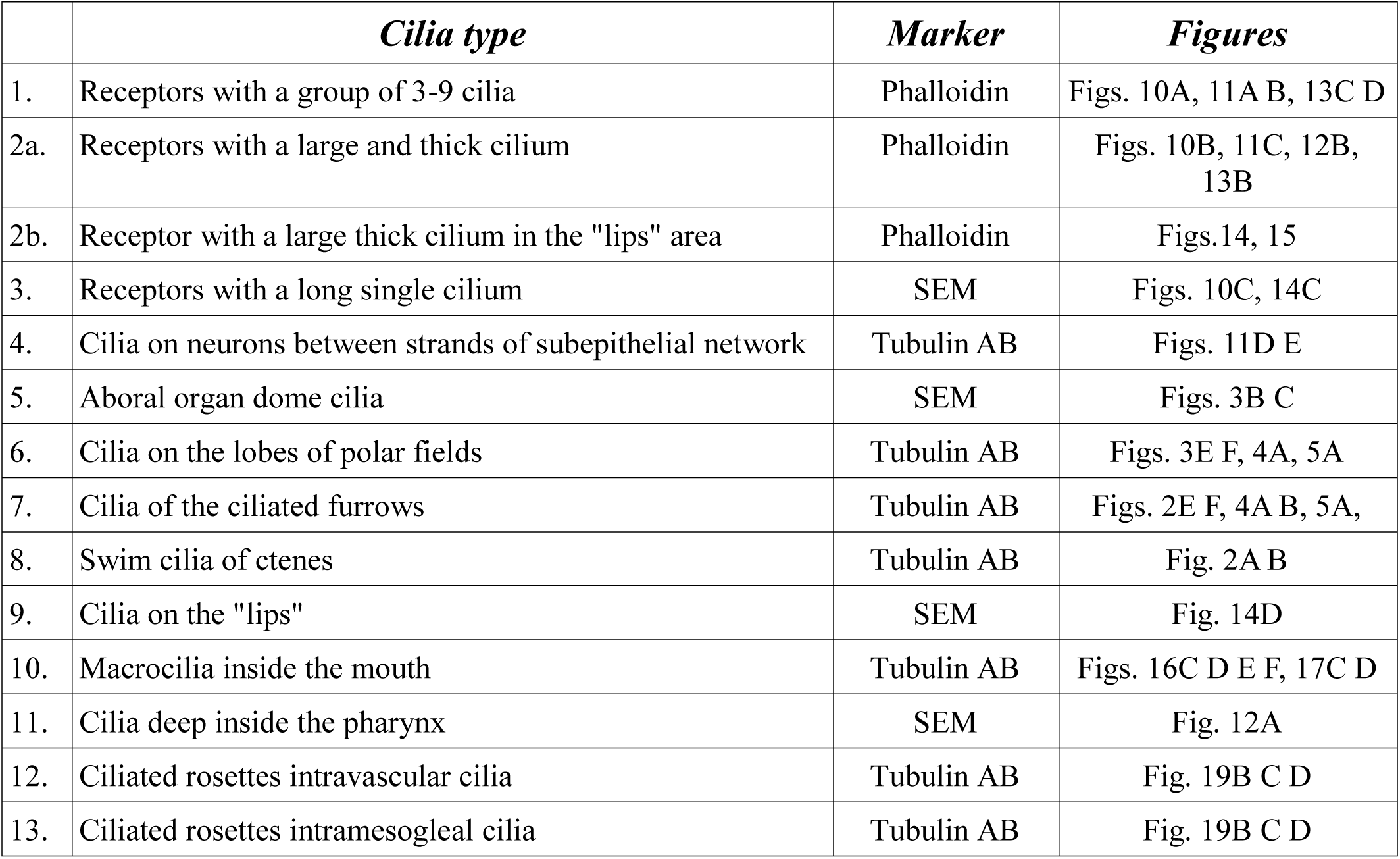
Types of cilia in *Beroe abyssicola*, and their markers.

**Table 4.** Muscle types in *Beroe* described in this study.

|  | *Muscle group* | *Figure* |
| --- | --- | --- |
| 1. | Large Circular muscles in mesoglea | Fig. 21 A B C |
| 2. | Large Longitudinal muscles in mesoglea | Fig. 21C |
| 3. | Radial muscles in mesoglea connecting the outside wall with the pharynx | Fig. 21C D |
| 4. | Long muscles attached to the comb rows | Figs. 21F |
| 5. | Non-contractile muscle-like filaments in comb rows | Figs. 21F |

## 4. DISCUSSION

The deciphering ctenophore innovations has a great interest for biology and neuroscience (Striedter et al., 2014; Moroz, 2018; Sebe-Pedros et al., 2018) due to possible independent origins of many animal traits such as neurons, muscles, mesoderm, anus, gastrovascular and sensory systems as well as lack of reference cellular atlases for representatives of this basal metazoan phylum. Apparently complex evolutionary history of the phylum lead to substantial radiation within the ancestral body plan in ctenophores around the Permian, where the lineages leading to *Pleurobrachia* and *Beroe* were parted (Whelan et al., 2017). Combining each direction, around 500 million years separate cydippids and non-tentacular ctenophores with remarkable distinct morphologies and life-styles. However, as we have found, the basic structure of the nervous system is very similar among ctenophores: both subepithelial neural net and mesogleal network as well as comparable neuronal and receptor types do present in both studied groups. It implies substantial evolutionary conservative mechanisms to maintain the neuronal architecture across the ctenophore lineage.

This fact is confirmed by highly conservative organization of a unique “presynaptic triad” across ctenophore synapses (Hernandez-Nicaise, 1973c; Hernandez-Nicaise, 1991). Nevertheless, except glutamate, ctenophores do not share any neurotransmitters with Bilateria and Cnidaria (Moroz et al., 2014; Moroz and Kohn, 2016). Most neurotransmitters in ctenophores are unknown, and the list of signal molecules include several dozen of the lineage-specific neuropeptides further supporting the convergent evolutions of neurons and synapses (Moroz et al., 2014; Moroz, 2015; Moroz and Kohn, 2016).

Nevertheless, within the same neuronal overall architecture - the same type of the polygonal subepithelial network (with predominantly bipolar and tripolar neurons) in both species, we observed differences derived from quite distinct morphologies and behaviors between *Beroe* and *Pleurobrachia* (Norekian and Moroz, 2018). Both *Pleurobrachia* and *Beroe* are predators. However, their hunting strategies are very different. *Pleurobrachia* spreads its tentacles and “fishes” for most of zooplankton, which can be caught in the net. *Beroe* is an active hunter swimming and searching for the prey, which demands acuter sensory perception. Below, we summarize some of lineage-specific traits and similarities.

First, the sizes of the polygonal units and neurons themselves are bigger in *Beroe* because of its superior size.

Second, *Pleurobrachia* has a pair of tentacles, which *Beroe* does not have. There are four tentacular nerves that control tentacle movements in *Pleurobrachia* (Norekian and Moroz, 2018), which are absent in *Beroe.* Interestingly, except for the aboral organ and polar fields, we did not detect any substantial concentration of neuronal elements or nerve analogs in *Beroe,* although we expected so due to its more complex swimming patterns and hunting behaviors.

Third, the subepithelial network in *Pleurobrachia* covers the entire wall of the tentacle pockets. *Beroe,* on the other hand, feeds on a large size prey, which is consumed in an enormous pharynx that extends throughout the entire body. As a result, the subepithelial neural network covers the entire surface of the pharynx and is very similar to the network in the skin, effectively doubling the cumulative size of the nerve net. We did not notice such network in much smaller *Pleurobrachia* pharynx (Jager et al., 2011; Norekian and Moroz, 2018), but further investigations are required.

Fourth, we identified the same three types of mesogleal neurons in both species: bipolar cells, multipolar cells, and cells giving rise to long thin processes crossing the mesoglea. One noticeable difference was that in *Pleurobrachia* multipolar, star-like neurons appeared to be most numerous, while in *Beroe* we observed relatively more bipolar neurons in the mesoglea compare to multipolar cells. The size of mesogleal neurons was also significantly larger in *Beroe,* but the density of cells was slightly higher in *Pleurobrachia*.

Fifth, we have identified the same five types of receptors associated with the diffused neural network in both species (Norekian and Moroz, 2018). The most numerous type on the surface were receptors with multiple cilia. In *Beroe*, these receptors were significantly bigger and more pronounced. At this stage, without functional data, it is difficult to speculate about the function and modality of ctenophore receptors. We hypothesize that most of single nonmotile cilium cells might be mechanoreceptors (number type 2a and 2b in Table 2). Very interesting receptors (type 5 in Table 2) with three long tubulin IR processes were located under the network itself, joining the strands of the polygonal network. They certainly never reached the surface, and probably represent proprioreceptors. Receptors with more flexible and/or numerous thin cilia might represent chemosensory elements (like type 3 in Table 2).

*Pleurobrachia* also have a unique receptor type located in the tentacles between colloblasts - *Beroe* does not have these receptors.

On the other hand, *Beroe* being a large active predator and a macro-feeder have a high density of receptors present on the oral side of the mouth in specialized sensory structures. The specialized sensory structures in *Beroe* included many sensory bulges, or bumbs, located on the surface just outside the mouth, which are densely covered by long mechanosensory cilia. Presumably, they are very important in providing the sensory information about the prey when *Beroe* swims with their mouth tightly closed. Also, *Beroe* have two types of receptors (with multiple cilia and a single thick cilium) present on the inside surface of the pharynx, albeit at much lower density than on the outer surface. It needs to be studied whether *Pleurobrachia* have similar receptors inside the pharynx.

Sixth significant difference is the presence of macrocilia in the mouth, which serve as teeth helping to hold the prey and push it down the pharynx, and even tear off pieces of tissue from a large prey. This feature is exclusive Beroids innovations (Horridge, 1965b; Tamm, 1982; Hernandez-Nicaise, 1991; Tamm, 2014), and *Pleurobrachia* does not have such unique cilia. Another specialized *Beroe* innovation is the presence of adhesive epithelial secretory cells in the “lips” area, which are instrumental in locking the lips together when *Beroe* is not feeding (Tamm and Tamm, 1991c). It appears to be an important function in the wide-mouthed *Beroe* during regular swimming activity. Those adhesive cells looked like elevated papillae in our SEM observations and showed distinct tubulin IR like other secretory cells in ctenophores.

Seventh, both the aboral organ and polar fields are the main sensory organs in ctenophores, and their structure and size find a reflection in their behavioral demands. Polar fields in *Pleurobrachia* are flat and narrow, while in *Beroe* they are very wide and have a large protruding “crown” of lobes and their branches covered with long cilia, which is elevated for about 200 µm above the epithelial layer and creates a more complex 3-dimensional structure.

Eight, there is a remarkable difference between these two species in the size, strength, and density of muscle fibers in their body. The body shape in *Pleurobrachia* is supported by rigid hydroskeleton, which significantly limits the movements of any body parts (other than tentacles) and does not require strong muscular support. In *Beroe,* on the other hand, the flexible and muscular body walls are instrumental in feeding and consuming the large prey in their huge pharynx. Therefore, *Pleurobrachia* have a loose mesh of external smooth parietal muscles just under their epithelium and neural network, plus relatively few thin muscle fibers are running throughout their mesoglea. Beroids do not have external parietal muscles (see Hernandez-Nicaise, 1991). However, they have thousands of truly gigantic muscles crossing the mesoglea, including circular, longitudinal and many radial muscles that connect outside integument with the wall of the pharynx. The differences in neuromuscular organization are also reflected in very strong defensive withdrawal responses, which can be seen in *Beroe* when the animal easily retracts the entire aboral organ and polar fields inside the body. *Pleurobrachia* does not have such powerful withdrawal responses. In contrast, it has a stereotyped “escape” rotational behavior, mainly controlled by the aboral organ. Although the overall comb rows structure is very conservative and similar between *Beroe* and *Pleurobrachia, Beroe* combs are bigger and swim cilia are significantly longer, but we did not find noticeable differences in comb innervation between species.

Ninth, the overall morphology of the gastrovascular system in *Beroe* and *Pleurobrachia* is also different. In *Beroe,* the system consists of meridional and pharyngeal canals with hundreds of branches and diverticula penetrating most of the mesoglea and acting as functional analogs of circulatory, hormonal, nutritional, excretory and reproductive systems. All branches and diverticula contain numerous pores - ciliated rosettes. These ciliated rosettes were also present on meridional canals of *Pleurobrachia*. Although, their basic structure was similar in both species, the ciliated rosettes were significantly bigger in *Beroe,* cilia were longer and were very brightly stained by tubulin antibody, while in *Pleurobrachia*, cilia had no tubulin IR and can be detected only by SEM.

**In summary**, in term of the ctenophore neural and receptor systems, we see more rearrangements within the same architecture, rather than extraordinary innovations. With the loss of tentacular systems in *Beroe,* we clearly detected the loss of tentacle nerves, but there is an expansion of surface nerve net neurons and receptors into the pharynx. The *Beroe* lineage has also developed the diversity of ciliated cells, including the ‘invention’ of the macrocilia, and evolved giant muscles with complex coordination mechanisms. About 40 distinct cell types were recognized in this study, which is essential for ongoing single-cell RNA-seq profiling in ctenophores and other reference species, critical to reconstruct the genealogy of neurons and other animal cell types (Moroz, 2018; Sebe-Pedros et al., 2018).

Combined, our data on two distantly related species revealed the unprecedented complexity of neuromuscular organization in this basal metazoan lineage with diverse types of neurons, receptors, ciliated cells and muscles to support ctenophore active swimming, complex escape and prey capture behaviors. The obtained neuroanatomy atlas provides unique examples of lineage-specific innovations within these enigmatic marine animals and reveals the remarkable architecture of sensory and effector systems.

## Acknowledgments

We thank FHL for their excellent collection and microscope facilities including the new Nikon Laser Scanning confocal microscope; Dr. Victoria Foe for the use of the BioRad confocal microscope and Dr. Adam Summers for the use of SEM. We also thank Dr. Claudia Mills and Dr. Billie Swalla for useful ctenophore discussions.

This work was supported by the United States National Aeronautics and Space Administration (grant NASA-NNX13AJ31G), the National Science Foundation (grants 1146575, 1557923, 1548121 and 1645219), and Human Frontiers Research Program and National Institute of Health (R01GM097502).

## Role of the authors

All authors had full access to all the data in the study and take responsibility for the integrity of the data and the accuracy of the data analysis. TPN and LLM share authorship equally. Research design: TPN, LLM. Acquisition of data: TPN, LLM. Analysis and interpretation of data: TPN, LLM. Drafting of the article: TPN, LLM. Funding: LLM.

